# Two-Stage CD8^+^ CAR T-Cell Differentiation in Patients with Large B-Cell Lymphoma

**DOI:** 10.1101/2025.03.05.641715

**Authors:** Guoshuai Cao, Yifei Hu, Tony Pan, Erting Tang, Nick Asby, Thomas Althaus, Jun Wan, Peter A. Riedell, Michael R. Bishop, Justin P. Kline, Jun Huang

**Affiliations:** Pritzker School of Molecular Engineering, University of Chicago, Chicago, IL 60637, USA; Pritzker School of Medicine, University of Chicago, Chicago, IL 60637, USA; Committee on Immunology, University of Chicago, Chicago, IL 60637, USA; Committee on Cancer Biology, University of Chicago, Chicago, IL 60637, USA; Department of Medicine, University of Chicago, Chicago, IL 60637, USA; The David and Etta Jonas Center for Cellular Therapy, University of Chicago, Chicago, IL 60637, USA; Department of Medical and Molecular Genetics, Indiana University School of Medicine, Indianapolis, IN 46202, USA

**Keywords:** large B-cell lymphoma, chimeric antigen receptor (CAR), axicabtagene ciloleucel, CD28-costimulated CAR, CAR T-cell differentiation, two-stage differentiation

## Abstract

Chimeric antigen receptor (CAR) T-cell therapy has expanded therapeutic options for patients with diffuse large B-cell lymphoma (DLBCL). However, progress in improving clinical outcomes has been limited by an incomplete understanding of CAR T-cell differentiation in patients. To comprehensively investigate CAR T-cell differentiation in vivo, we performed single-cell, multi-modal, and longitudinal analyses of CD28-costimulated CAR T cells from infusion product and peripheral blood (day 8-28) of patients with DLBCL who were successfully treated with axicabtagene ciloleucel. Here, we show that CD8^+^ CAR T cells undergo two distinct waves of clonal expansion. The first wave is dominated by CAR T cells with an exhausted-like effector memory phenotype during the peak expansion period (day 8-14). The second wave is dominated by CAR T cells with a terminal effector phenotype during the post-peak persistence period (day 21-28). Importantly, the two waves have distinct ontogeny and are biologically uncoupled. Furthermore, lineage tracing analysis via each CAR T cell’s endogenous TCR clonotype demonstrates that the two waves originate from different effector precursors in the infusion product. Precursors of the first wave exhibit more effector-like signatures, whereas precursors of the second wave exhibit more stem-like signatures. These findings suggest that pre-infusion heterogeneity mediates the two waves of in vivo clonal expansion. Our findings provide evidence against the intuitive idea that the post-peak contraction in CAR abundance is solely apoptosis or extravasation of short-lived CAR T cells from peak expansion. Rather, our findings demonstrate that CAR T-cell expansion and persistence are mediated by clonally, phenotypically, and ontogenically distinct CAR T-cell populations that serve complementary clinical purposes.

## INTRODUCTION

Diffuse large B-cell lymphoma (DLBCL), the most common non-Hodgkin’s lymphoma in the United States, is characterized by diffusely proliferating and malignant B cells at nodal or extranodal sites.^1^ Although up-front chemoimmunotherapy is often curative, patients with DLBCL that is refractory to up-front treatment or relapse following remission (r/r DLBCL) have limited treatment options and poor outcomes.^2,3^ Effective treatment options were limited until the United States Food and Drug Administration approved autologous CD19-directed chimeric antigen receptor (CAR) T-cell therapies for r/r DLBCL in 2017. Autologous CD19-directed CAR T-cell therapy involves virally transducing a patient’s T cells ex vivo with a CD19-directed CAR—an engineered receptor consisting of an extracellular anti-CD19 single-chain variable fragment, a hinge/transmembrane region, and intracellular costimulatory (CD28 or 4-1BB) and activation (CD3ζ) domains. CAR-transduced T cells (i.e., CAR T cells) are cultured and infused into the patient, where they lyse CD19^+^ lymphoma cells.^4^ CD19-directed CAR T-cell therapy has achieved complete response rates of 40-54%^5–8^ for r/r DLBCL, but non-response rates^6,9^ and treatment toxicities^10^ remain as challenges.^11^

Development of CAR T-cell formulations with higher response rates and fewer toxicities requires a thorough understanding of how CAR T cells differentiate in patients with r/r DLBCL. The pioneering ZUMA-1 trial (NCT02348216) for axicabtagene ciloleucel (autologous CD28-costimulated CAR T cells) demonstrated that peripheral blood CAR T cells expand, contract, and sometimes persist.^5^ Greater expansion predicted higher response rates, but also higher likelihood of developing treatment toxicities.^5,12^ Longer persistence predicted durable remission and long-term immunosurveillance with leukemias, but its relevance for preventing DLBCL relapse remains obscure.^9,13^ Although many factors are associated with expansion and persistence (including CAR design^14,15^ and CAR T-cell phenotypes^16–20^, among others^21–23)^, how and why CAR T cells differentiate into expansive or persistent phenotypes in vivo is still an open question. Addressing this question requires a single-cell approach that integrates CAR T-cell phenotypes and clonal kinetics over longitudinal timepoints. However, existing studies have either focused on CAR T-cell phenotypes^19,24,25^ or clonal kinetics^26^ without integrating both data modalities, or lacked the temporal resolution required to decipher cell fates over longer periods.^12,27^ Consequently, a complete and longitudinal understanding of CAR T-cell differentiation in vivo has remained elusive.

To comprehensively elucidate CAR T-cell differentiation in vivo, we performed single-cell, multi-modal (paired RNA-seq/CITE-seq/TCR-seq), and longitudinal analyses of CD28-costimulated CAR T cells from infusion product and peripheral blood of seven patients with r/r DLBCL who were complete responders under treatment with axicabtagene ciloleucel. Peripheral blood CAR T cells were sorted using CD19 antigen-tetramers.^28^ Importantly, we report that the CD8^+^ CAR T cells observed during the peak expansion period (day 8-14) have distinct clonotypic repertoires compared to the later CD8^+^ CAR T cells observed during the post-peak persistence period (day 21-28). This discovery led us to analyze how these two CAR T-cell populations differ with respect to phenotypes, transcriptional profiles, regulatory networks, and infusion product precursors. Our findings not only offer a finer understanding of CAR T-cell biology in vivo, but also inform efforts to develop CAR T cells with improved expansion and persistence.

## RESULTS

### Study design and clinical findings

To interrogate CD28-costimulated CAR T-cell differentiation in vivo, we longitudinally interrogated the phenotypes and clonal dynamics of CAR T cells from seven patients (P1-7) who achieved complete responses under treatment with axicabtagene ciloleucel (**Figure 1a**). Patients were diagnosed with r/r DLBCL and treated at the University of Chicago Medicine between 2019 and 2021 (table S1). Clinical response was determined by positron emission tomography/computed tomography imaging 30 days after infusion product administration. Complete response was defined as no detectable lymphoma (figure S1). To capture phenotypic heterogeneity and clonal dynamics longitudinally, we performed single-cell multi-modal analyses (paired RNA-seq/CITE-seq/TCR-seq via the 10x Genomics platform) on infusion product and peripheral blood biospecimens at three timepoints: peak expansion (T_exp_, day 8-14), early post-peak persistence (T_per1_, day 21), and late post-peak persistence (T_per2_, day 28) (table S2). The three peripheral blood timepoints were available for all patients, except P3 (only T_exp_ and T_per1_). CAR T cells were sorted from peripheral blood using CD19 antigen-tetramers (representative staining in **Figure 1b**, figure S2).^28^ Compared to CD19 antigen-tetramer-negative T cells, CD19 antigen-tetramer-positive T cells specifically expressed the CAR transgene, which validates our sorting strategy (**Figure 1c**).

**Figure 1.**
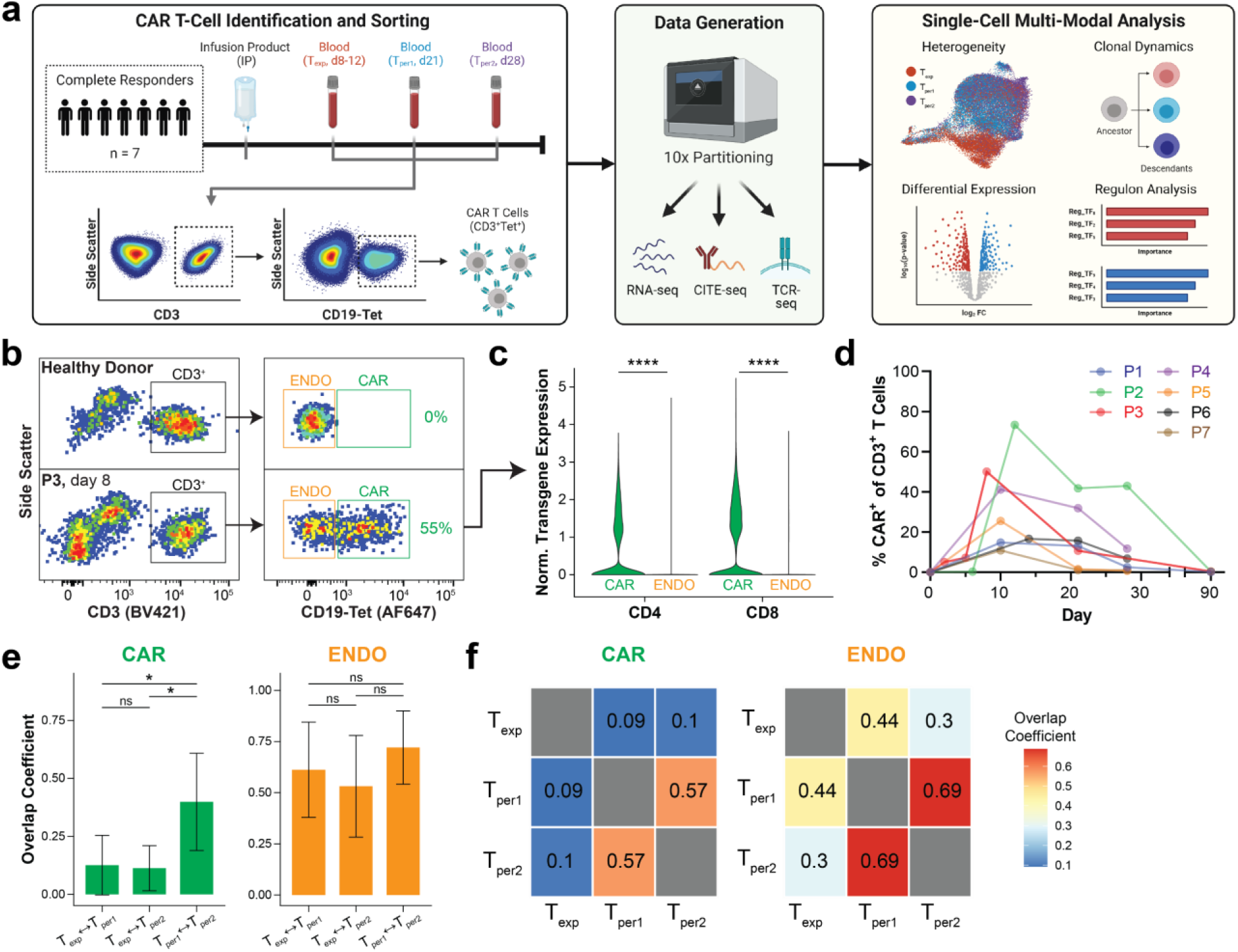
CD8^+^ CAR T cells from complete responders undergo a clonotypic shift in vivo. (**a**) Schematic depicting sorting strategy, data generation, and single-cell multi-modal analysis of CAR T cells in peripheral blood from seven CAR T-cell therapy patients (P1-7) who exhibited complete responses with axicabtagene ciloleucel. (**b**) Representative flow plots depicting anti-CD3 and CD19 antigen-tetramer staining of peripheral blood mononuclear cells from P3 versus a healthy donor. CD3^+^Tet^+^ (“CAR”) and CD3^+^Tet^−^ (“ENDO”) patient cells were sorted from the indicated gates. (**c**) Violin plots depicting normalized CAR transgene mRNA expression of sorted CAR and ENDO T cells, split by CD4^+^ (left) and CD8^+^ (right) subsets. Expression levels were compared by Wilcoxon Rank-Sum test, whereby **** indicates p<0.0001. (**d**) Line plots depicting expansion and contraction of peripheral blood CAR abundance over the course of therapy. (**e-f**) Bar graphs and heatmaps depicting overlap coefficients for TCR clonotypes comparing CAR (left) and ENDO (right) repertoires between T_exp_, T_per1_, and T_per2_. Overlap coefficients were compared by *t*-test, whereby * indicates p<0.05 and ns indicates not significant.

### CD8^+^ CAR T cells undergo a clonotypic shift between T_exp_ and T_per_

To analyze CAR T-cell population dynamics, we tracked CAR abundance (% CAR^+^ of CD3^+^ T cells) in peripheral blood throughout the course of therapy. CAR abundance at peak expansion ranged from 11% to 73% (**Figure 1d**). Peak expansion occurred at day 8-14, which is consistent with prior clinical findings.^5^ For each timepoint, we quantified proportions of CD8^+^ and CD4^+^ T cells within the total CAR T-cell population by single-cell RNA-seq and CITE-seq. CAR T cells were predominantly CD8^+^ across most patients and timepoints (figure S3).

We next analyzed the dynamics of the CAR T-cell clonal repertoire across longitudinal timepoints (T_exp_ at day 8-14, T_per1_ at day 21, T_per2_ at day 28) using single-cell TCR-seq. Clone sizes were similar across timepoints (figure S4a). Clonotypes did not overlap between patients (figure S4b). Repertoire overlap analysis indicated that T_exp_ clonotypes were significantly distinct from T_per1_ or T_per2_ clonotypes (**Figure 1e-f**, left). In sharp contrast, T_per1_ and T_per2_ clonotypes overlapped substantially more. As a control, we also analyzed endogenous (non-CAR) T cells from matched timepoints. Unlike with CAR T cells, endogenous T cells did not show distinctive clonotypic patterns (**Figure 1e-f**, right), indicating that the shift in clonotypes between T_exp_ and T_per_ is CAR-specific. This CAR-specific clonotypic shift was consistent across patients (figure S4b). Moreover, the distinction between T_exp_ and T_per_ clonotypes is driven by CD8^+^ T cells, and not by CD4^+^ T cells (figure S4c). Collectively, these findings indicate that CD8^+^ CAR T cells undergo a clonotypic shift between T_exp_ and T_per_.

### CD8^+^ CAR T cells undergo a phenotypic shift from exhausted-like EM to TE

Having shown a shift in CD8^+^ CAR T-cell clonotypes between T_exp_ and T_per_, we hypothesized that unique CD8^+^ CAR T-cell phenotypes dominate T_exp_ and T_per_. To test this hypothesis, we filtered CD8^+^ CAR T cells for Uniform Manifold Approximation and Projection (UMAP) and identified six T-cell clusters (**Figure 2a**) based on gene and protein markers (**Figure 2b**, expanded marker set in figure S5a). No cluster was patient-specific (figure S5b). All clusters expressed CAR transgene and CD8α, validating our sorting and filtering processes, respectively. All clusters expressed *CXCR3*, a chemokine receptor that demarks activated T cells.

**Figure 2.**
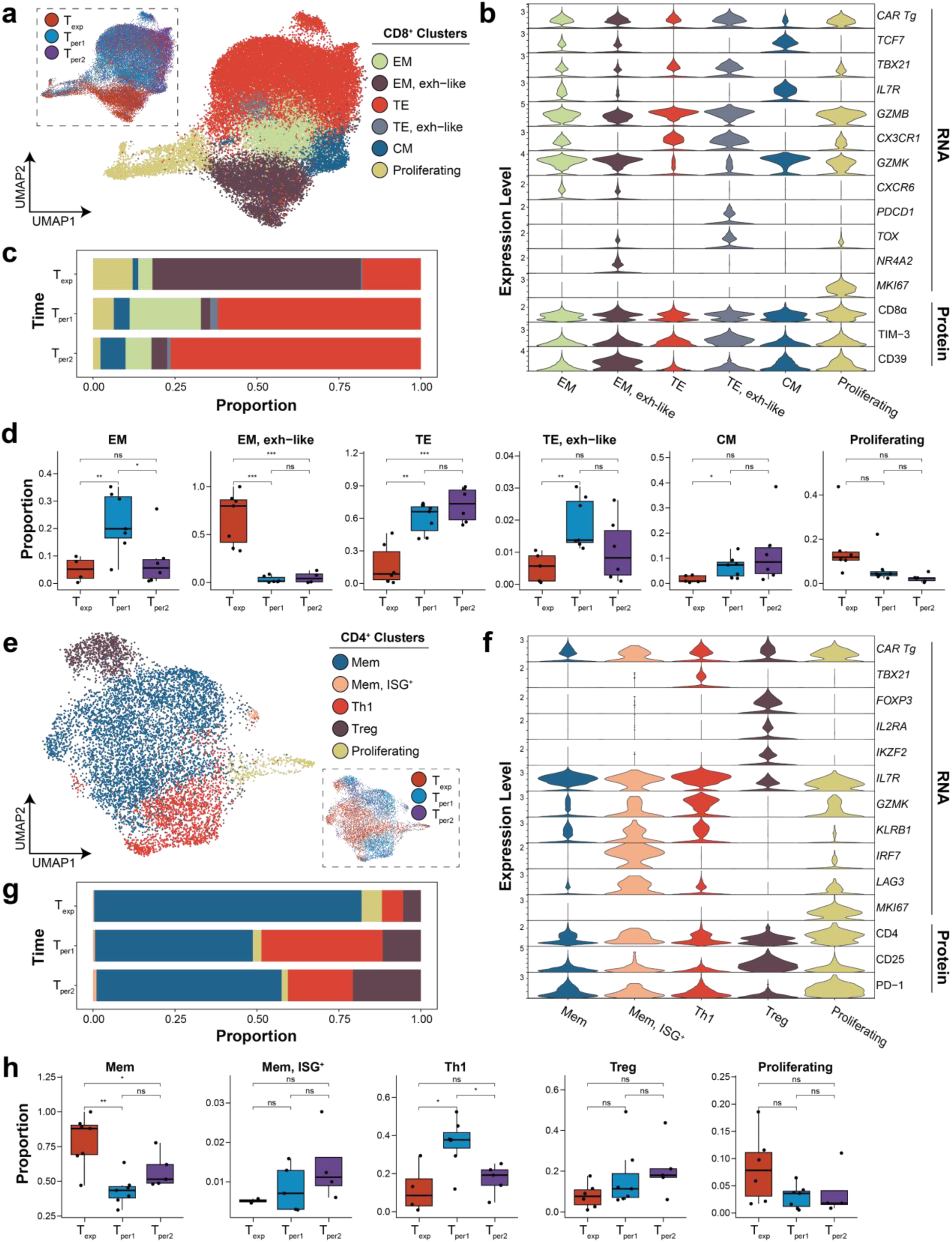
Phenotypic heterogeneity of peripheral blood CD8^+^ and CD4^+^ CAR T cells. (**a**, **e**) UMAPs depicting single-cell transcriptomes of CD8^+^ (**a**) and CD4^+^ (**e**) CAR T cells colored by cell cluster. Inset depicts distribution of transcriptomes across timepoints. (**b**, **f**) Violin plots depicting normalized expression levels of key genes and proteins for annotating and phenotyping CD8^+^ (**b**) and CD4^+^ (**f**) CAR T cells. For extended versions, see figure S5a and S6a. (**c**, **g**) Stacked bar graphs depicting proportions of each CD8^+^ (**c**) and CD4^+^ (**g**) CAR T-cell phenotype at different timepoints. (**d**, **h**) Boxplots depicting proportion of CD8^+^ (**d**) and CD4^+^ (**h**) CAR T cells of a given phenotype at different timepoints. Each dot represents a measurement from a single patient. Proportions are compared between timepoints by Wilcoxon Rank-Sum test, whereby *** indicates p<0.001, ** indicates p<0.01, * indicates p<0.05, and ns indicates not significant. Abbreviations: CM, central memory; EM, effector memory; TE, terminal effector; Mem, memory; ISG, interferon stimulated genes; Th1, type 1 helper-like

Examination of gene and protein markers revealed that one of the six clusters represented proliferating T cells (*MKI67*^+^*TOP2A*^+^). The remaining five non-proliferating clusters were annotated as central memory (CM, *TCF7*^+^*TBX21*^-^), effector memory (EM, *TCF7*^+^*TBX21*^+^), or terminal effector (TE, *TCF7*^-^*TBX21*^+^) T cells. The *TCF7*^+^*TBX21*^-^ CM cluster exhibited markers of stemness (*IL7R*, high CD127) and minimal markers of effectorness (*GZMB*, *CX3CR1*). The two *TCF7*^+^*TBX21*^+^ EM clusters occupied the lower half of the UMAP and uniquely expressed *CXCR6*, a chemokine receptor that facilitates trafficking into solid tumors.^29^ One of these EM clusters upregulated markers consistent with early exhaustion (*NR4A2*, *TOX*, *GZMK*, low TIM-3), hence it was designated “exhausted-like EM”. Lastly, the two *TCF7*^-^*TBX21*^+^ TE clusters occupied the upper half of the UMAP and uniquely downregulated *GZMK*. One of these TE clusters upregulated markers consistent with late exhaustion (*TOX*, *PDCD1*, high TIM-3), hence it was designated “exhausted-like TE”. The other TE cluster was highly clonal (some clone sizes > 100), suggesting expansion through proliferation (figure S7a). Overall, most CD8^+^ CAR T cells were *TBX21*^+^ EM or TE, which is consistent with the established link between CD28 costimulation and effector memory (rather than central memory) differentiation.^14,15^

Phenotypic compositions of CD8^+^ CAR T cells at T_exp_ and T_per_ were compared. CAR T cells at T_exp_ were predominantly exhausted-like EM (64%) whereas CAR T cells at T_per_ were predominantly TE (63% for T_per1_, 77% for T_per2_) (**Figure 2c**). These findings were statistically significant and consistent across all seven patients (**Figure 2d**, figure S5c). Moreover, the large clone sizes within the T_per_-specific TE cluster (figure S7a) suggest active TE proliferation at T_per_. CAR T cells at T_per1_ were enriched for EM. From T_per1_ to T_per2_, the EM proportion decreased while the TE proportion increased, suggesting progressive differentiation from EM to TE. However, the overall phenotypic compositions at T_per1_ and T_per2_ were more similar than different, which is concordant with findings from repertoire overlap analysis (**Figure 1e-f**). Moreover, we observed decreasing proliferating proportions and increasing CM proportions over time (**Figure 2d**), though this was not always statistically significant. These changing proportions may suggest some CAR T cells were returning from an activated to a resting phenotype. In conclusion, CD8^+^ CAR T cells phenotypically shifted from exhausted-like EM to TE between T_exp_ and T_per_.

### CD4^+^ CAR T cells maintain a memory phenotype with CAR Treg persistence

We next performed clustering, annotation, and longitudinal analyses of CD4^+^ CAR T cells to investigate phenotypes at T_exp_ and T_per_. We identified five T-cell clusters, all expressing CAR transgene and CD4, based on gene and protein markers (**Figure 2e-f**; expanded marker set in figure S6a). No cluster was patient-specific (figure S6b). All clusters expressed *CXCR3*, indicating T-cell activation and type 1 helper polarization.

One of the five clusters represented proliferating T cells (*MKI67*^+^*TOP2A*^+^). The remaining four non-proliferative clusters were annotated as memory (Mem, *TBX21*^-^*FOXP3*^-^), type 1 helper (Th1, *TBX21*^+^*FOXP3*^-^), or regulatory (Treg, *TBX21*^-^*FOXP3*^+^) T cells. The two *TBX21*^-^*FOXP3*^-^ Mem clusters at the center of the UMAP comprised most of the cells. One of the Mem clusters upregulated *IRF7* and interferon-stimulated genes (*MX1*, *OAS1*, *ISG15*), indicating response to type I interferon signaling. This signature suggests dynamic interferon secretion in vivo and is in concordance with type I interferon’s role in memory CD4^+^ T-cell differentiation.^30^ The *TBX21*^+^*FOXP3*^-^ Th1 cluster along the lower half of the UMAP upregulated cytolytic genes (*PRF1*, *GNLY*, *GZMK*, *NKG7*) and tissue-homing chemokine receptors (*CX3CR1*, *CXCR6*), resembling the cytotoxic CD4^+^ CAR T cells described by Melenhorst et al.^19^ Lastly, the *TBX21*^-^*FOXP3*^+^ Treg cluster expressed classic Treg markers (*IL2RA*, CD25, low *IL7R*, low CD127) and upregulated *IKZF2*, indicating a suppressive phenotype.^31^ CAR Tregs inhibit conventional CAR T cells^27,32^ and portend progressive disease.^12,27^

Phenotypic compositions of CD4^+^ CAR T cells at T_exp_ and T_per_ were compared. CD4^+^ CAR T cells predominantly exhibited a Mem phenotype (49-82%) across all timepoints (**Figure 2g**). This observation was consistent across all patients (figure S6c). Between T_exp_ and T_per1_, the Mem proportion significantly decreased while the Th1 proportion significantly increased, indicating that CD4^+^ CAR T cells may be increasingly polarized during the early contraction period (**Figure 2h**). Moreover, across all patients, the Treg proportion steadily increased from T_exp_ (5%) to T_per1_ (11%) to T_per2_ (21%), indicating that CAR Tregs were maintained in peripheral blood after peak expansion. The CAR Treg cluster was predominantly non-clonal (figure S7b), suggesting CAR Tregs were maintained through persistence rather than proliferation. While previous studies have only investigated CAR Tregs at day 7^12,27^, this current study affirms and extends the presence of CAR Tregs until at least day 28. Importantly, persistence of CAR Tregs points towards their possible involvement in decreasing acute inflammation and restoring immune homeostasis after peak expansion. In conclusion, we discovered that CD4^+^ CAR T cells did not exhibit an abrupt T_exp_-to-T_per_ phenotypic shift. Rather, they consistently exhibited a memory phenotype with CAR Tregs persisting over the course of therapy.

### Integration of clonotypic and phenotypic shifts in CD8^+^ CAR T cells supports a two-stage differentiation model

Having observed shifts in both CD8^+^ CAR T-cell clonotypes and phenotypes between T_exp_ and T_per_, we hypothesized the existence of two distinct waves of in vivo clonal expansion. To test this hypothesis, we linked the clonotype and phenotype of single CD8^+^ CAR T cells in our dataset using their cell barcodes as unique indices. After this linking process, we confirmed that the proportions of T cells with exhausted-like EM (EM-exh) and terminal effector (TE) phenotypes (**Figure 3a**) are consistent with prior results (**Figure 2c**). A phenotype label was subsequently assigned to each CD8^+^ CAR T-cell clone based on its predominant phenotype at each timepoint. For each patient, we tracked the total abundances of clones sharing a common phenotype label across timepoints. Notably, the abundance of EM-exh clones at T_exp_ was significantly reduced at T_per1_ and T_per2_ (**Figure 3b**, left), while the abundance of TE clones at T_per1_ and T_per2_ was significantly reduced at T_exp_ (**Figure 3b**, middle and right). In addition to redemonstrating the shifts in CD8^+^ CAR T-cell clonotypes and phenotypes, these findings support the existence of two distinct waves of in vivo clonal expansion. (**Figure 3c**).

**Figure 3.**
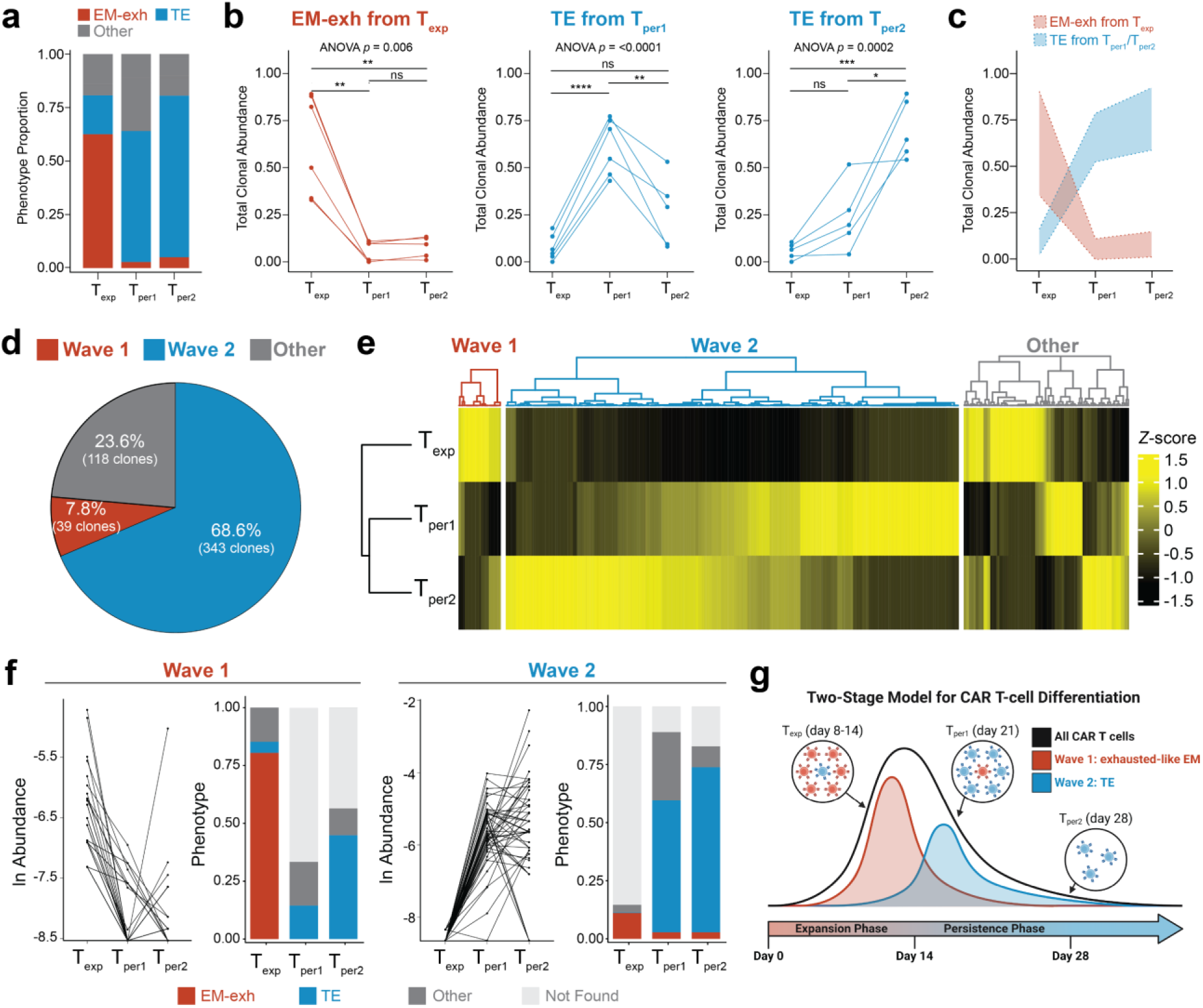
Integration of clonotypic and phenotypic shifts supports a two-stage differentiation model. All figure panels are based on the clonotype-phenotype linked dataset (n=32432 cells). (**a**) Proportion of CAR T cells with an exhausted-like effector memory (EM-exh), terminal effector (TE), or other phenotypes at each timepoint. (**b**) Total abundance of all clones that were predominantly EM-exh at T_exp_ (left), TE at T_per1_ (middle), or TE at T_per2_ (right), measured across timepoints. A clone’s predominant phenotype was defined as the phenotype with the greatest representation. Each point represents clones from a patient (n=6). Abundance across timepoints were compared by repeated measures ANOVA with post-hoc *t*-tests. (**c**) Overlay of clonal abundance dynamics with 95% confidence intervals for clones annotated as EM-exh from T_exp_ (red) or TE from T_per1_/T_per2_ (blue). (**d**) Pie chart depicting proportion of the top 500 clones classified as Wave 1, Wave 2, or Other. (**e**) Heatmap depicting normalized CAR abundance across timepoints for the top 500 largest clones. (**f**) Clonal dynamics and phenotypic distribution for the top 500 clones, grouped into Wave 1 and Wave 2. Left panels depict the change in clonal abundance across timepoints, while right panels show the predominant phenotype distribution at each timepoint. (**g**) Cartoon summarizing the two-stage model for CAR T-cell differentiation. Bulk CAR T-cell expansion and contraction (black line) masks the dynamics of Wave 1 (EM-exh, expansion phase timeframe, red) and Wave 2 (TE, persistence phase timeframe, blue) clones.

Next, we evaluated the explanatory power of our hypothesis for two distinct clonal waves. To minimize noise, we filtered for the top 500 largest clones (mean size 10.8, minimum size 4).

Resulting clones were annotated with the following three definitions, based on our hypothesis for two waves: “Wave 1” (phenotypically EM-exh with highest abundance at T_exp_), “Wave 2” (phenotypically TE with highest abundance at T_per1_/T_per2_), or “Other” (not Wave 1 or Wave 2). Wave 1 and Wave 2 clones together accounted for a significant majority (76%) of the top 500 clones (**Figure 3d**). In accordance with our definitions, Wave 1 clones were maximally abundant at T_exp_, while Wave 2 clones were maximally abundant at T_per_ (**Figure 3e**). Wave 1 clones predominantly exhibited an EM-exh phenotype at T_exp_ and sharply decreased in abundance at T_per_ (**Figure 3f**, left). In contrast, Wave 2 clones predominantly exhibited a TE phenotype at T_per_ and sharply decreased in abundance at T_exp_ (**Figure 3f**, right). To investigate whether these findings are dependent on clone size, we expanded our analysis to the top 3000 largest clones (mean size 3.7, minimum size 2). Under this filter, Wave 1 and Wave 2 clones together continued to account for a significant majority (79%) of the top 3000 clones (figure S8a) and reaffirm the expected clonal and phenotypic dynamics (figure S8b-c). Therefore, our hypothesis for two distinct waves of in vivo clonal expansion has high explanatory power and is robust to clone size.

In conclusion, the two distinct waves of in vivo clonal expansion strongly substantiate a two-stage CD8^+^ CAR T-cell differentiation model (**Figure 3g**). Under this model, some CAR T cells (Wave 1) with an exhausted-like EM phenotype expand earlier to dominate the peak expansion timeframe (T_exp_), while other CAR T cells (Wave 2) with a TE phenotype expand later to dominate the post-peak persistence timeframe (T_per_). Of note, the two-stage differentiation model uncouples CD8^+^ CAR T cells from peak expansion and post-peak persistence by designating these two waves as separate lineages. Moreover, these findings provide evidence against the intuitive idea that the post-peak contraction in CAR abundance is solely apoptosis or extravasation of short-lived CAR T cells from peak expansion. Rather, even as total CAR abundance contracts after peak expansion, a distinct subset of CAR T-cell clones simultaneously expands to eventually dominate the peripheral blood CAR T-cell repertoire.

### T_exp_-and T_per_-specific transcriptional signatures and regulatory networks

We set out to identify the molecular determinants underlying CD8^+^ CAR T cells at T_exp_ and T_per_ using gene set enrichment analysis (GSEA), differential gene expression analysis (DGEA), and regulatory network analysis. For internal validation, the average expression of timepoint-specific genes (see Methods) for each sample (patient by timepoint) were hierarchically clustered and compared via correlation (**Figure 4a**). Samples largely clustered by timepoint, validating the existence of patient-independent molecular signatures. We then validated our dataset against external data from Maus et al., which consisted of CD8^+^ CAR T cells from patients with large B-cell lymphoma (**Figure 4a-b**, colored in tan and brown, six in total after filtering).^27^ Based on their sample collection timing (day 7), external data from Maus et al. should resemble our samples at T_exp_, rather than at T_per_. Consistent with expectations, external data resembled T_exp_ transcriptomes on both pseudo-bulk (**Figure 4a**, via correlation) and single-cell (**Figure 4b**, via label transfer) levels, which increases the external validity of our findings.

**Figure 4.**
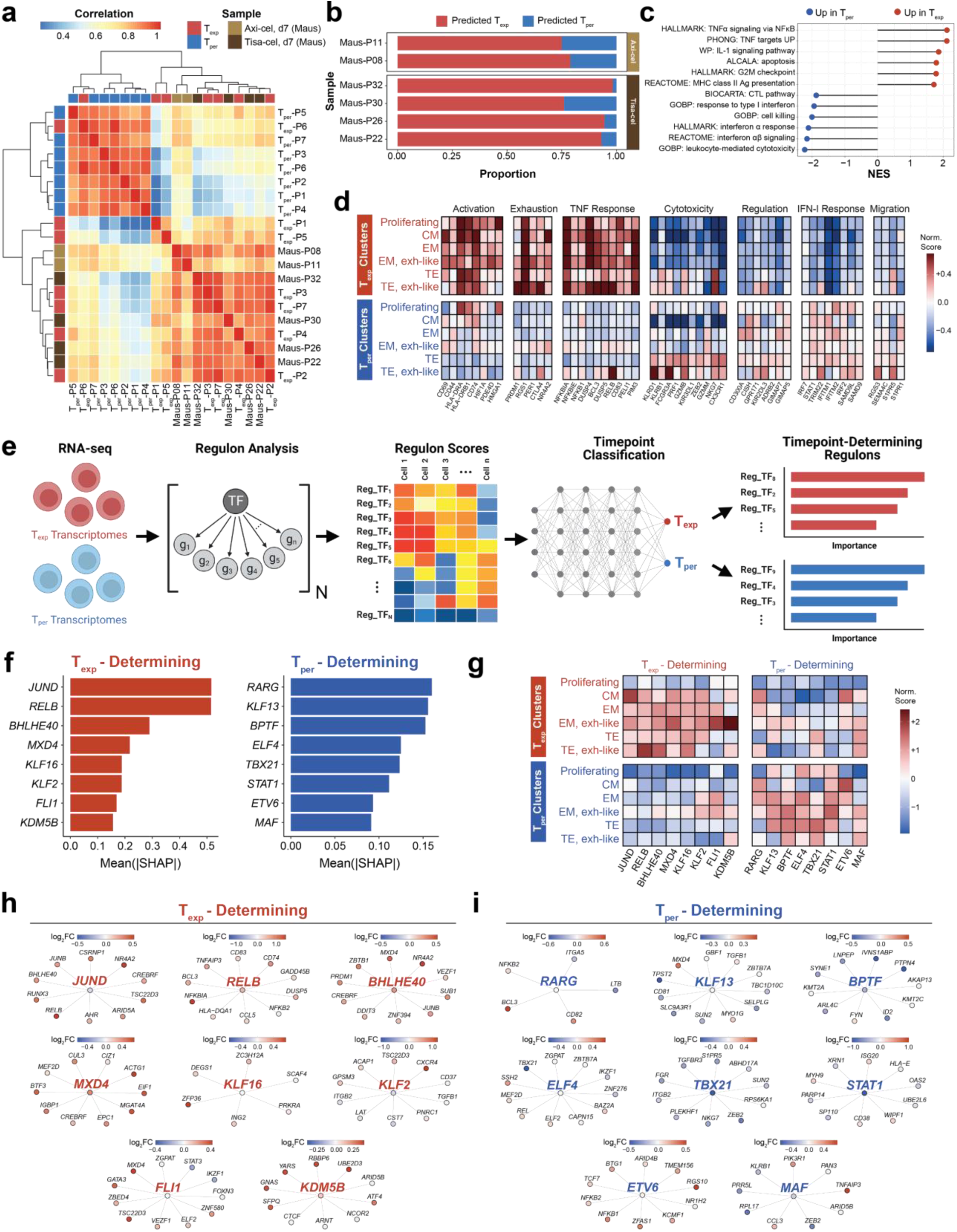
Transcriptional signatures and regulatory networks of CD8^+^ CAR T cells at T_exp_ and T_per_. (**a**) Heatmap depicting correlation between pseudo-bulk transcriptome of each sample (patient by timepoint). Samples were ordered along columns and rows by hierarchical clustering. Transcriptomes of day 7 samples from Maus et al. were added for external validation. (**b**) Stacked bar graph depicting label transfer of timepoint (T_exp_ or T_per_) from this study’s dataset onto single-cell transcriptomes of samples from Maus et al. (**c**) Gene set enrichment analysis comparing CAR T cells between T_exp_ and T_per_. Gene sets were ordered by direction of upregulation and magnitude of enrichment. (**d**) Tile map depicting normalized expression of genes (columns) among different cell clusters at T_exp_ and T_per_ (rows). Genes were manually grouped into modules according to known functions. (**e**) Schematic for regulon construction and T_exp_ versus T_per_ classification. After transcriptomes were transformed into regulomes, regulon scores were calculated to train a machine-learning model to classify CAR T cells from T_exp_ and T_per_. Key regulons were identified based on importance for the model’s predictions. (**f**) Bar graph depicting the SHapley Additive exPlanation (SHAP) values for the top eight regulons underlying T_exp_ and T_per_ predictions. (**g**) Tile map depicting normalized signature scores of regulons (columns) among different cell clusters at T_exp_ and T_per_ (rows). Regulons were grouped as T_exp_-determining (left) and T_per_-determining (right). (**h**, **i**) Target networks for the top eight T_exp_-determining (**h**) and T_per_-determining regulons (**i**). In each regulatory network, only the top differentially expressed genes are depicted. Each gene is colored according to log_2_ fold-change between expression in T_exp_ (red) and T_per_ (blue).

To elucidate the immunological processes at each timepoint, CD8^+^ CAR T-cell transcriptomes at T_exp_ and T_per_ from our seven patients were compared via GSEA (**Figure 4c** and table S3 for single-cell method, figure S9a and table S4 for pseudo-bulk method) and DGEA (**Figure 4d**, figure S9b, tables S5-6). CAR T cells at T_exp_ upregulated gene sets for cell cycling and apoptosis, consistent with a short-lived phenotype. Upregulation of the cell cycling gene set remained true even when only comparing the proliferating cell cluster at T_exp_ and T_per_ (figure S9d). This indicates that, while proliferating cells are present at both T_exp_ and T_per_, those at T_exp_ are more proliferative than those at T_per_. We also observed T_exp_-specific and cluster-independent upregulation of genes related to T-cell activation (including *CD69*, *CD44*, *CD74*) and exhaustion (including *PRDM1*, *CTLA4*, *NR4A2*), implicating antigen-engagement and CAR-mediated signaling during T_exp_. On the other hand, CAR T cells at T_per_ upregulated gene sets for cytotoxicity and immune regulation. Expression of cytotoxicity-related genes (including *KLRB1*, *FCGR3A*, *PRF1*, *GZMB*, *NKG7*) was restricted to TE and exhausted-like TE clusters, indicating that this gene signature arises from T_per_-specific preponderance of the terminal effector phenotype. We also observed T_per_-specific upregulation of regulatory genes, including GTPase immune-associated proteins (*GIMAP5*, *GIMAP7*) which predict long-term persistence^33^, as well as sphingosine-1-phosphate receptors (*S1PR1*, *S1PR5*) which indicate blood localization^34^. These signatures suggest that CAR T cells at T_per_ are functional and capable of long-term persistence in peripheral blood. Lastly, downregulation of genes related to T-cell activation and exhaustion implicate decreased in vivo antigen load and/or CAR-mediated signaling during T_per_.

Interestingly, some differentially expressed gene signatures between CD8^+^ CAR T cells at T_exp_ and T_per_ pertained to cytokine signaling (**Figure 4c-d**, see figure S9c). CAR T cells at T_exp_ upregulated TNF response genes (including *NFKBIA*, *NFKBIE*, *DUSP4*, *RELB*). TNF can be secreted by CAR T cells for autocrine signaling^20^, predicts efficacious CAR T cells^20^, and contributes to CRS.^35^ On the other hand, CAR T cells at T_per_ upregulated type I interferon (IFN-I) response genes (including *STAT1*, *IFITM1*, *IFITM2*, *IRF7*). Many IFN-I response genes have known antiviral functions. The temporally specific and cluster-independent upregulation of TNF and IFN-I response genes indicates a dynamic in vivo cytokine environment during the CAR T-cell immune response.

Next, we investigated T_exp_-and T_per_-specific regulatory networks using a machine-learning model (**Figure 4e**, see Methods). Single-cell transcriptomes were transformed into regulomes to calculate regulon scores. A machine-learning model was trained using the regulon scores to classify CD8^+^ CAR T cells at T_exp_ and T_per_. This strategy resulted in high (∼90%) classification accuracy across patients (figure S10a). Top T_exp_-and T_per_-determining regulons were identified based on each regulon’s importance (SHAP value) for the model’s prediction (**Figure 4f**, table S7). Sizes of timepoint-determining regulons span approximately 10–1000 genes (figure S10b). Regulon expression was compared between cell clusters and timepoints (**Figure 4g**). T_exp_-and T_per_-determining regulons were broadly upregulated at T_exp_ and T_per_, respectively. However, expression of some regulons was cluster-specific (e.g., *FLI1* regulon among EM clusters).

The top three T_exp_-determining regulons (for *JUND*, *RELB*, *BHLHE40*) exhibited higher expression among non-proliferating clusters and included co-regulated inflammation-associated genes (including *NFKBIA*, *TNFAIP3*, *DUSP5*) (**Figure 4h**, see figure S11a).^37^ The *RELB* and *BHLHE40* genes themselves are included in the *JUND* regulon, indicating they may be directly upregulated by *JUND*. These three regulons were upregulated at T_exp_ and suggest response to pro-inflammatory cytokines (such as TNF, discussed above) or CAR signaling. *BHLHE40* rewires mitochondrial metabolism for tissue residency^38^, which may contribute to CAR T-cell fitness after entering the lymphoma. Shared genes between the *JUND* and *BHLHE40* regulons included AP-1 transcription factors (*JUN*, *JUNB*, *FOSL2*), which have been implicated in T-cell and CAR T-cell exhaustion^39,40^, and may contribute to the exhausted-like EM phenotype that predominates the first clonal wave at T_exp_. Among remaining T_exp_-determining regulons, *FLI1* is notable for antagonizing effector T-cell differentiation.^41^ Consistent with known biology, expression of the *FLI1* regulon was upregulated among EM clusters, downregulated among TE clusters, and did not significantly change between T_exp_ and T_per_ within TE clusters.

On the other hand, T_per_-determining regulons were more obscure (**Figure 4i**, see figure S11b). Among top T_per_-determining regulons, *KLF13* regulates T-cell apoptosis^42^, *BPTF* activates a stemness gene expression program^43^ and maintains peripheral T-cell homeostasis^44^, and *ELF4* promotes memory CD8^+^ T cells by regulating quiescence^45^. Altogether, the putative functions of the *KLF13*, *BPTF*, and *ELF4* regulons may underlie the delayed expansion and persistence of the second clonal wave at T_per_. Moreover, these functions indicate restoration of immune homeostasis during T_per_. The *TBX21* regulon was notable for effector-associated genes (including *NKG7*, *ZEB2*) and was upregulated within non-CM clusters, consistent with preponderance of the TE phenotype at T_per_. The *TBX21* regulon also includes *S1PR5*, which is a known T-BET target^46^ and retains T cells in peripheral blood^34^. Lastly, the *STAT1* regulon was notable for interferon-associated antiviral genes (including *OAS2*, *ISG20*), in concordance with upregulation of IFN-I signaling gene sets at T_per_.

### CD8^+^ CAR T cells at T_exp_ and T_per_ originate from distinct precursors in the infusion product

The distinct clonotypes, phenotypes, transcriptional signatures, and regulatory networks underlying the two waves of CD8^+^ CAR T cells led us to hypothesize that they originate from distinct precursors in the infusion product. To test this hypothesis, we first investigated infusion product phenotypes. After filtering for infusion product CD8^+^ T cells, we identified four clusters based on gene expression density (**Figure 5a-b**, figure S12b) and levels (**Figure 5c**, figure S12a): proliferating (high *MKI67*), activated naïve-like (naïve-act, high *TCF7*), type 1 effector (EFF-Tc1, high *TBX21*), and type 2 effector (EFF-Tc2, high *GATA3*) T cells. The *TCF7*^hi^ naïve-act cluster upregulated markers of naiveness (*IL7R*, *LEF1*, *CCR7*, CD45RA) and T-cell activation (*CD38*, *CD95*), consistent with an early activated or stem cell-like memory phenotype. The two EFF clusters upregulated cytolytic molecules (*GZMB*, *PRF1*, *GNLY*) and downregulated markers of naiveness (*IL7R*, *LEF1*). EFF-Tc1 and EFF-Tc2 were distinguished by expression density of lineage-specific transcription factors (*TBX21* for Tc1, *GATA3* for Tc2) and receptors (*KLRG1* for Tc1, *CCR4* for Tc2). The EFF-Tc1 cluster exhibited lower CAR transgene expression than the EFF-Tc2 cluster (figure S12b-c), which may impact their functional phenotypes in vivo.^47,48^ The three non-proliferating clusters upregulated *IRF7* and anti-viral genes (*ISG20*, *IFITM1*, *IFITM2*), indicating type I interferon signaling during ex vivo transduction and/or expansion. Cluster proportions varied between patients (figure S12d). In general, proliferating T cells (54%) were the most prevalent, while proportions of non-proliferating phenotypes (naïve-act, 14%; EFF-Tc1, 11%; EFF-Tc2, 21%) were mutually comparable (**Figure 5e**, top row).

**Figure 5.**
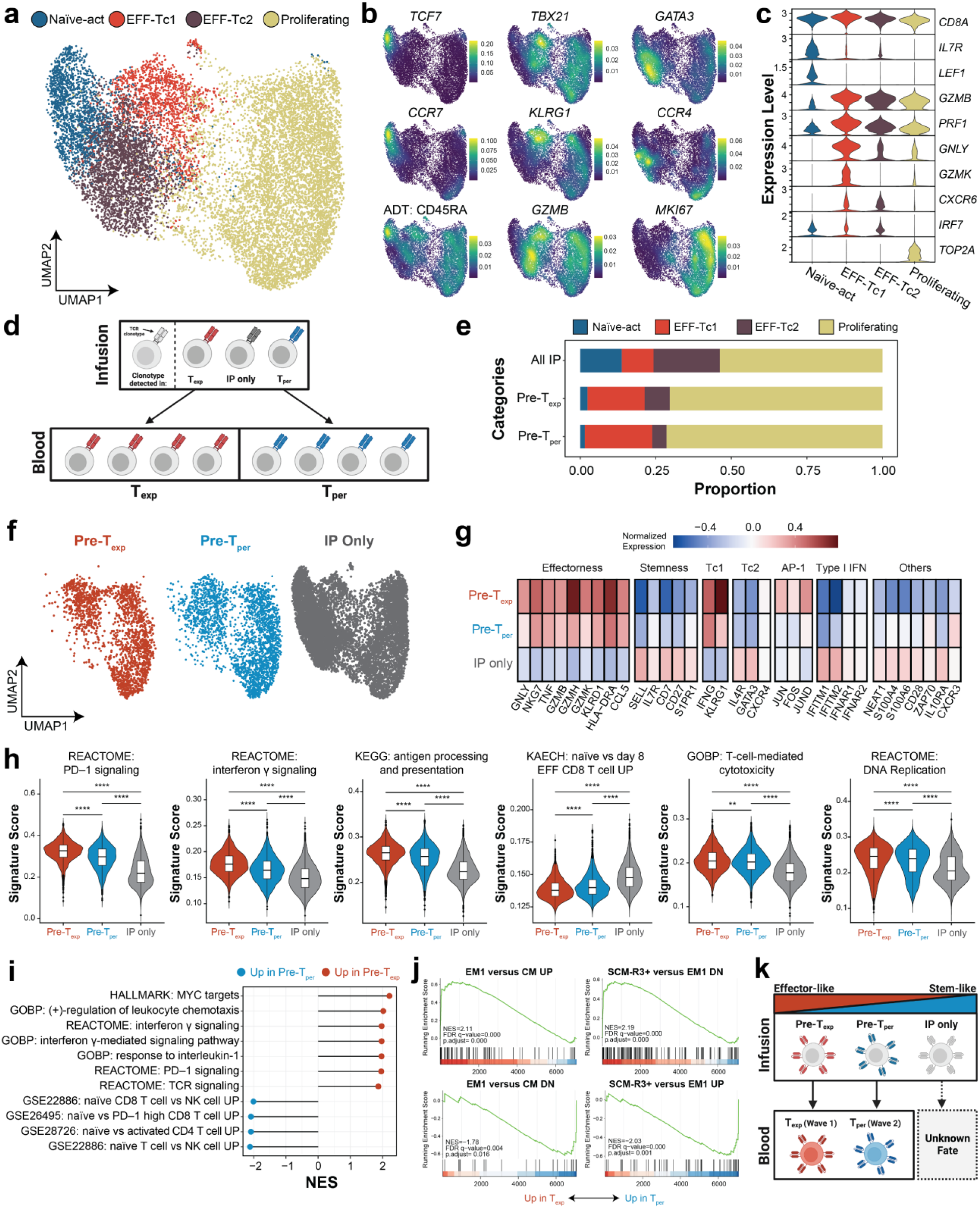
Infusion product precursors of peripheral blood CD8^+^ CAR T cells. (**a**) UMAP depicting single-cell transcriptomes of infusion product CD8^+^ T cells colored by cell cluster. (**b**, **c**) Density maps (**b**) and violin plots (**c**) depicting expression levels of key genes and proteins for annotation and phenotyping. For extended version, see figure S11a-b. (**d**) Cartoon depicting identification of Pre-T_exp_ and Pre-T_per_ using endogenous TCR clonotypes as unique indices. (**e**) Stacked bar graph depicting proportion distribution of all infusion product (top row) or precursors of peripheral blood CD8^+^ CAR T cells (bottom two rows) among the four cell clusters. (**f**) Colored UMAPs depicting distribution of Pre-T_exp_, Pre-T_per_, and non-linked infusion product cells (“IP only”) on the overall UMAP. (**g**) Tile map depicting normalized expression of genes (columns) among Pre-T_exp_, Pre-T_per_, and IP only infusion product cells (rows). Genes were manually grouped into modules according to known functions. (**h**) Violin plots depicting expression levels of select differentially expressed gene sets between Pre-T_exp_, Pre-T_per_, and IP only infusion product cells. Expression levels were compared by Wilcoxon Rank-Sum test, whereby **** indicates p<0.0001 and ** indicates p<0.01. (**i**) Gene set enrichment analysis comparing Pre-T_exp_ and Pre-T_per_. Gene sets were ordered according to direction of upregulation and magnitude of enrichment. (**j**) Enrichment plots for select gene sets differentially expressed between Pre-T_exp_ and Pre-T_per_ infusion product cells. (**k**) Cartoon depicting fates of CD8^+^ CAR T cells over the entire course of therapy, from infusion product precursors to peripheral blood CAR T cells at T_exp_ and T_per_. Abbreviations: NES, normalized enrichment score

Having established infusion product phenotypes, we next performed TCR lineage tracing analysis to developmentally link infusion product CAR T cells with peripheral blood CAR T cells using their endogenous TCR clonotypes as unique indices. This method delineates infusion product precursors of CD8^+^ CAR T cells at T_exp_ (“Pre-T_exp_”) and at T_per_ (“Pre-T_per_”) (**Figure 5d**). Both Pre-T_exp_ and Pre-T_per_ comprised all four infusion product clusters (**Figure 5e**, bottom rows) and largely overlapped on the UMAP (**Figure 5f**). Hence, we conclude that coarse cluster phenotype alone cannot accurately predict in vivo differentiation. Compared to T cells not linked to peripheral blood (“IP only”), Pre-T_exp_ and Pre-T_per_ cells upregulated expression of effectorness (including *GZMB*, *TNF*, *GNLY*) and type 1 polarization (including *IFNG*, *KLRG1*) genes, as well as lower expression of stemness (including *SELL*, *IL7R*) and type 2 polarization (including *GATA3*, *IL4R*) genes (**Figure 5g**). The significance of type 1 versus type 2 polarized CD8^+^ CAR T cells is not well-understood in the literature and warrants future studies. Pathway analysis indicated that Pre-T_exp_ and Pre-T_per_ upregulated gene sets for T-cell effector function (“PD-1 signaling”, “T-cell-mediated cytotoxicity”), response to interferon γ (“interferon γ signaling”, “antigen processing and presentation”), and DNA replication, whereas non-linked CAR T cells upregulated gene sets for stemness (**Figure 5h**, table S8). Our discovery of CD8^+^ CAR T-cell precursors with an effector-like phenotype is corroborated by findings from Thomas et al., who also identified effector-like precursors from CAR T-cell patients treated for B-cell acute lymphoblastic leukemia.^49^ Possible fates for the non-linked, naïve-like, IP-only CAR T cells include extravasation into the lymphoma, expansion in peripheral blood outside the T_exp_-T_per_ window (day 8-28), or failure to persist in vivo.

Lastly, we directly compared transcriptomic signatures of Pre-T_exp_ and Pre-T_per_ via GSEA. Although both Pre-T_exp_ and Pre-T_per_ broadly exhibited an effector phenotype (**Figure 5g-h**), Pre-T_exp_ upregulated effectorness-associated gene sets (including TCR, PD-1, interferon γ, and IL-1 signaling), whereas Pre-T_per_ upregulated stemness-associated gene sets (**Figure 5i**). In concordance, DGEA showed that Pre-T_exp_ upregulated effector molecules (*GZMB*, *GZMK*, *FCGR3A*, *NKG7*, *KLRG1*), AP-1 transcription factors (*JUN*, *JUND*, *FOS*), and MHC class II expression (*HLA-DRB1*, *HLA-DRB5*, *HLA-DRA*), whereas Pre-T_per_ upregulated naïve-like markers (*SELL*, *IL7R*, *S1PR1*) (figure S12e, table S9). Upregulation of *JUN* in Pre-T_exp_ confers exhaustion resistance^39^, potentially underlying their eventual efficacy at T_exp_. Interestingly, Pre-T_per_ exhibited greater CAR transgene expression (figure S12e), which may reflect transduction differences among apheresis precursors or a link between CAR expression and in vivo differentiation.^48^ Next, we compared Pre-T_exp_ and Pre-T_per_ using transcriptomic signatures from a human CD8^+^ differentiation atlas from Giles et al.^50^ Pre-T_exp_ more closely resembled an effector memory state, whereas Pre-T_per_ more closely resembled either a stem-cell memory or central memory state (**Figure 5j**). Altogether, these patterns indicate that, while both Pre-T_exp_ and Pre-T_per_ exhibited an effector phenotype in the infusion product, Pre-T_exp_ were more differentiated with greater effectorness, whereas Pre-T_per_ were less differentiated with greater stemness.

In conclusion, TCR lineage tracing analysis supports our hypothesis that CD8^+^ CAR T cells at T_exp_ and T_per_ originate from different infusion product precursors. Integrating these findings into the two-stage differentiation model paints a more complete picture of CD8^+^ CAR T cells over the month following CAR T-cell administration (**Figure 5k**). Effector CD8^+^ CAR T cells exist along a gradient of effectorness and stemness in the infusion product. Following infusion into the patient, effector CAR T cells with greater effectorness rapidly expand until peak expansion (T_exp_, days 8-12), adopting a functional and cytotoxic EM phenotype with exhaustion-like characteristics upon antigen stimulation. Lymphoma infiltration and in vivo killing^51^, CRS^5^, and ICANS^5^ coincide with the T_exp_ timeframe, suggesting that these CAR T cells mediate tumor clearance and side effects. Subsequently, this first wave of expanded CAR T cells diminishes through apoptosis or extravasation. Simultaneously, the remaining effector CAR T cells with greater stemness from the infusion product expand during the post-peak persistence timeframe (T_per_, day 21-28). These newer and longer-lived CAR T cells adopt effector characteristics and persist in vivo, where they may ensure a durable response through long-term immunosurveillance.

### CD8^+^ Exhausted-like EM CAR T cells at T_exp_ exhibit characteristics of early exhaustion

Discovery of the predominant exhausted-like EM phenotype in peripheral blood at T_exp_ may be surprising since exhaustion was conventionally defined in lymphoid organs or tumor-infiltrating lymphocytes.^52^ To verify this annotation, we further characterized the molecular signatures of the exhausted-like EM cluster. The exhausted-like EM cluster was densely situated at the lower half of the UMAP (**Figure 6a-b**). We used RNA-seq/CITE-seq to examine gene and protein expression density, respectively (**Figure 6c**, figure S13a for additional markers). The exhausted-like EM cluster highly expressed exhaustion-associated transcription factors (*NR4A2*, *TOX*, *IRF4*) and inhibitory receptors (*ENTPD1*, *PDCD1*, *TIGIT*, *LAG3*, *CTLA4*). Intermediate expression of memory (*TCF7*, *LEF1*, *CD27*, *IL7R*) and effector (*TBX21*, *GZMB*, *PRF1*, *GNLY*, *IFNG*, *NKG7*) genes was also observed. Additionally, the cluster exhibited low expression of *CX3CR1* (effector lineage marker) and *B3GAT1* (senescence marker). CITE-seq measurements of protein expression for cell surface receptors (including CD39, PD-1, TIGIT, LAG3, CD27, CD57, CD127, TIM-3, CXCR3) were largely consistent with corresponding gene expression. Sequencing data indicated that high expression of CD39 (protein for *ENTPD1*) and low expression of CD57 (protein for *B3GAT1*) most clearly differentiated the exhausted-like EM cluster from other clusters. To validate these characteristics, flow cytometry was employed to analyze a set of longitudinal patient PBMCs. Consistent with sequencing data, flow cytometry data demonstrated that the CD39^+^CD57^-^ phenotype was highest among CD8^+^ CAR T cells at T_exp_ (75%) compared to at T_per1_ (33%) and T_per2_ (18%) (**Figure 6d**, top row). Furthermore, flow cytometry showed decreased CX3CR1 expression at T_exp_, and increased expression at T_per1_ and T_per2_, in agreement with sequencing data (**Figure 6d**, bottom row). This mixed expression pattern of exhaustion, memory, and effector markers resembles the circulating PD1^+^CD39^+^ T_ex_ cells described by the Wherry group’s human T-cell differentiation atlas.^50^

**Figure 6.**
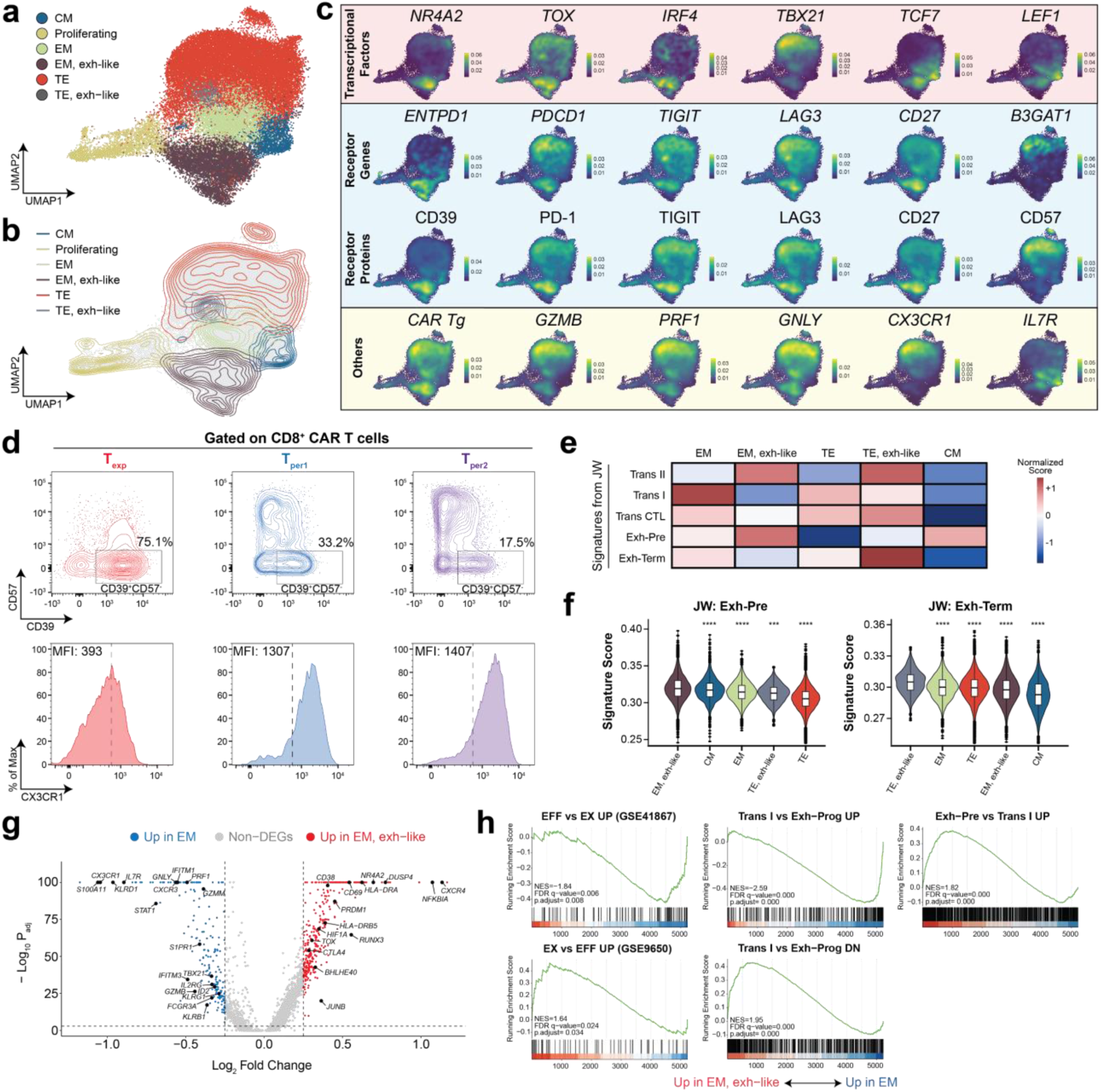
Transcriptomic signatures of exhausted-like EM CD8^+^ CAR T cells. (**a**, **b**) UMAP depicting single-cell transcriptomes of peripheral blood CD8^+^ T cells colored by cell cluster (**a**) or density contours of each cluster (**b**). The exhausted-like EM cluster is located on the lower half of the UMAP. (**c**) Density maps depicting expression levels of major T-cell genes and proteins, divided into categories. In the “receptor” category, proteins are placed directly beneath the corresponding gene. (**d**) Flow plots for validating transcriptomic data. Plots depict expression of CD39 and CD57 (top row) or CX3CR1 (bottom row) in CD8^+^ CAR T cells at each timepoint from P2. Exhausted-like EM CAR T cells were expected to be T_exp_-specific with high CD39 and low CD57/CX3CR1 expression. (**e**, **f**) Heat map depicting normalized expression of major gene sets from Wherry et al. (**e**). Expression of two of the gene sets were depicted as violin plots (**f**), ordered by decreasing expression level per cluster. Expression levels were compared to that of the cluster with highest expression via Wilcoxon Rank-Sum test, with p-values adjusted for multiple hypotheses testing using the Benjamini-Hochberg method, whereby **** indicates p<0.0001, *** indicates p<0.001. (**g**) Volcano plot depicting differentially expressed genes between EM and exhausted-like EM CD8^+^ CAR T cells. Genes were colored according to direction of upregulation. (**h**) Enrichment plots for select gene sets differentially expressed between EM and exhausted-like EM CD8^+^ CAR T cells.

Next, we conducted GSEA using gene sets from Wherry et al. that identified T-cell differentiation states in the Armstrong/clone 13 LCMV model (**Figure 6e**).^53^ These gene sets constitute comprehensive references for defining T-cell exhaustion because they (1) incorporate standing knowledge from the entire field, (2) integrate single-cell RNA-seq and ATAC-seq data from serial timepoints, and (3) originate from the Wherry group where comprehensive transcriptional signatures of T-cell exhaustion and most exhausted subsets were first defined. The two EM clusters (EM and exhausted-like EM) in our dataset correlated with different states from the Wherry group’s model. Our EM cluster resembled the “transitional I” T cells from the Armstrong model, indicating that these cells are not exhausted. Conversely, our exhausted-like EM cluster resembled the “precursor exhausted” T cells from the clone 13 model (**Figure 6f**, left). Simultaneously, the exhausted-like EM cluster did not resemble the “terminally exhausted” T cells from the same model (**Figure 6f**, right), which instead correlated with our exhausted-like TE cluster. Given clear differences between the EM and exhausted-like EM clusters, we directly compared their transcriptomic signatures through DGEA and GSEA. The exhausted-like EM cluster upregulated genes associated with T-cell activation (*CD69*, *CD38*, *DUSP4*, *NFKBIA*) and exhaustion (*TOX*, *NR4A2*, *PRDM1*, *CTLA4*) (**Figure 6g**, table S10). In contrast, the EM cluster upregulated genes associated with effector function (*TBX21*, *CX3CR1*, *GZMB*, *PRF1*, *FCGR3A*), memory (*IL7R*, *S1PR1*), and type I interferon signaling (*STAT1*, *IFITM1*, *IFITM3*). GSEA revealed that the exhausted-like EM cluster exhibited more exhaustion signatures and fewer effector signatures compared to the EM cluster (**Figure 6h**, left column, see also figure S13b). GSEA also confirmed that the exhausted-like EM cluster resembled the “exhausted progenitor” and “exhausted precursor” subsets from Wherry et al. (whereas the EM cluster resembled the “transitional I” subset) (**Figure 6h**, middle and right columns). Our findings suggest that the exhausted-like EM cluster exhibits robust gene and protein signatures of early exhaustion (justifying the “exhausted-like” annotation), yet is also phenotypically distinct from effector memory T cells.

## DISCUSSION

In this study, we use a combination of single-cell RNA-seq/CITE-seq/TCR-seq and longitudinal analyses to investigate CD28-costimulated CAR T-cell differentiation in seven patients with r/r DLBCL. Our findings are summarized in our two-stage model for CAR T-cell differentiation. Specifically, CD8^+^ CAR T cells undergo two distinct clonal expansion waves (at peak expansion and post-peak persistence timeframes), as revealed by clonotypic, phenotypic, and linked clonotypic-phenotypic analyses. The two waves are dominated by exhausted-like effector memory and terminal effector phenotypes, respectively. The exhausted-like effector memory annotation is supported by a CD39^+^CD57^-^ flow phenotype, low CX3CR1 expression, and an early exhaustion signature. We also identify transcription factors and regulatory networks associated with the first wave (including *JUND*, *RELB*, *BHLHE40*, *FLI1* regulons) and second wave (including *KLF13*, *BPTF*, *ELF4*, *TBX1*, *STAT1* regulons). Lastly, lineage tracing analysis determined that CD8^+^ CAR T cells from both waves derive from effector precursors in the infusion product. However, effector precursors of the first wave exhibit more effector-like signatures, whereas effector precursors of the second wave exhibit more stem-like signatures, suggesting that pre-infusion heterogeneity mediates two-stage differentiation. Our two-stage model implies that manipulating the phenotypic composition of the infusion product may allow more precise control over in vivo CAR T-cell differentiation for modulating therapeutic efficacy.

Although two-stage differentiation is a novel phenomenon that has not yet been reported in the literature, the individual elements of our model are consistent with current understanding in the field. Previous studies on 4-1BB-costimulated CAR T-cell differentiation described bursts of CAR T-cell clonal expansion that lead to significant changes in the CAR T-cell clonal repertoire over time.^19,54^ Our study not only generalizes this observation to CD28-costimulated CARs, but also directly pairs changes in the clonal repertoire with changes in T-cell phenotypes. Furthermore, previous reports have described CAR T-cell clones with memory-like phenotypes that exhibit delayed expansion after infusion^23^, which corroborates with the second clonal expansion wave at T_per_. Finally, the T_exp_-and T_per_-specific phenotypes described in our model are consistent with prior phenotyping studies.^25,27,49,55,56^ We also report agreement between our transcriptomes at T_exp_ and external data from Maus et al.^27^ By integrating our findings from multiple data modalities with established studies in the CAR T-cell field, we present a more complete and coherent two-stage model for in vivo CD28-costimulated CAR T-cell differentiation.

The two-stage differentiation model we present in this study has an important implication: CD28-costimulated CD8^+^ CAR T cells from the peak expansion and post-peak persistence timeframes are biologically uncoupled. This uncoupling significantly informs how we understand CAR T-cell expansion and persistence. Expansion facilitates rapid tumor clearance but can also cause CRS and ICANS^5,17^, while persistence facilitates long-term immunosurveillance but can also cause B-cell aplasia and hypogammaglobulinemia.^9,13,57^ Our findings suggest that expansion and persistence, both of which serve complementary clinical purposes, are mediated by distinct CD8^+^ CAR T-cell populations. Effective and personalized CAR T-cell therapies balance expansion and persistence, taking into consideration each patient’s tumor burden, tolerance for side effects, and risk of relapse. Our findings suggest that expansion and persistence may be independently tuned to meet a patient’s needs. On the other hand, our findings also suggest that engineering CD8^+^ CAR T cells that simultaneously expand and persist competently may be challenging, given that these characteristics originate from uncoupled CAR T-cell populations in vivo.

In addition to clonal kinetics and phenotypic heterogeneity, we characterized T_exp_-and T_per_-specific upregulation of TNF and IFN-I response gene sets and regulons, respectively. Notably, TNF-secreting CD8^+^ CAR T cells have been associated with complete responders^20^, suggesting that TNF may be signaling in an autocrine or paracrine manner during T_exp_. Upregulation of IFN-I response genes during T_per_ was intriguing. IFN-I response signatures in the apheresis of patients with B-cell acute lymphoblastic leukemia predict poor CAR T-cell persistence^58^, but IFN-I signaling may also enhance CAR T-cell efficacy in vivo.^59^ Complicating this story further, we also identified IFN-I response signatures in the infusion product, which implies that interferons are secreted by an unknown source during ex vivo transduction and/or expansion. Hence, the possible roles of IFN-I in CAR T-cell differentiation are likely complex, context-dependent, or time-dependent, and warrant further mechanistic investigation.

We concluded our study by using single-cell TCR-seq to link CD8^+^ CAR T cells at T_exp_ and T_per_ with effector precursors in the infusion product that exhibit more effector-like or more stem-like signatures, respectively. Discovery of effector precursors is externally corroborated by findings from Thomas et al^49^. Importantly, this linkage suggests that the magnitude or duration of peak expansion and post-peak persistence can be modulated by manipulating the relative quantities of precursors in the infusion product. However, we cannot rule out the possibility that the in vivo peripheral blood environment in a patient with DLBCL also plays an instrumental role in determining whether a CAR T cell clonally expands at T_exp_ or T_per_. Future studies can investigate this possibility by analyzing interactions between CAR T cells and other blood cells (such as myeloid cells, B cells, or NK cells) at T_exp_ and T_per_.

The present study focuses on CD28-costimulated CAR T cells from complete responders, but two-stage differentiation may be more widely applicable. Others forms of adoptive cell therapy, such as 4-1BB-costimulated CAR T cells for B-cell leukemias^60^ or systemic lupus erythematosus^61^, may also exhibit a two-stage differentiation pattern in vivo. Indeed, our transcriptomic signatures at T_exp_ were highly consistent with external data from day 7 tisagenlecleucel CD8^+^ CAR T cells from Maus et al.^27^, hinting that two-stage differentiation may be more general. Different forms of adoptive cell therapy and disease contexts may exhibit different temporal dynamics or favor one wave over the other. Two-stage differentiation may also be aberrant in non-responders. Future studies are needed to explore potential differences in temporal dynamics, cell phenotypes, or major regulatory pathways at longitudinal timepoints in alternative therapeutic contexts.

Although this current study represents an advance for the CAR T-cell therapy field, we acknowledge four limitations. Firstly, we analyzed CAR T cells in two locations: peripheral blood and infusion product. However, CAR T cells are also found elsewhere, including lymph nodes and lymphoma foci. Importantly, CAR T-cell differentiation may be influenced by interactions with antigen-presenting cells in lymph nodes or with tumor cells in the lymphoma. Our study was ultimately limited by sample availability. Future studies can focus on less accessible locations and determine CAR T cell fates beyond peripheral blood. Secondly, we studied CAR T-cell intrinsic factors (RNAs, proteins, clonotypes) that can influence differentiation. However, differentiation may also depend on interactions between CAR T cells and a complex in vivo environment that includes other leukocytes (myeloid cells, B cells, NK cells), lymphoma cells, and cytokines. Thirdly, we focused on CAR T cells longitudinally between peak expansion and post-peak persistence timeframes. Hence, we cannot rule out additional waves of clonal expansion in the near (˂1 week after infusion) or long (>1 month after infusion) terms. Lastly, we lack knowledge on whether two-stage differentiation applies to other contexts, such as non-responders, 4-1BB-costimulated CAR T cells, or TCR-transduced T cells. Future studies can explore alternative clinical contexts and test the generality of the two-stage differentiation model.

## METHODS

### Patient biospecimen collection

Deidentified biospecimens were obtained from a cellular therapy biobank in accordance with the institutional review board at the University of Chicago Medicine. Ethical guidelines were followed.

Longitudinal peripheral blood mononuclear cells (PBMCs) were collected from peripheral blood biospecimens by Ficoll-Paque PLUS (Cytiva, 95021-205) and stored in freezing media (RPMI, 10% FBS, 10% DMSO). Residual infusion product cells were collected from the patient’s spent infusion product bag and stored in CELLBANKER 1 (Amsbio, 11888). Cells were cryopreserved in liquid-phase nitrogen.

### Generation of CD19-tetramers

Tetramers^28^ were constructed from AviTag-biotinylated human His-tagged CD19 (Acro Biosystems, CD9-H82E9) and Alexa Fluor 647-labeled streptavidin (BioLegend, 405237). Biotinylated CD19 was added to the tetrameric streptavidin at a 4:1 molar ratio for 30 minutes at 4°C in the dark. This mixture was diluted with PBS to convenient concentrations for staining. For each batch of biotinylated CD19, quality controls were performed by generating CD19-tetramers and quantifying their ability to stain anti-CD19 CAR-transduced Jurkat T cells.

### Single-cell RNA-seq/CITE-seq/TCR-seq

Cryopreserved biospecimens were thawed (RPMI, 10% FBS) and washed with cold FACS buffer (PBS, 2% BSA, 0.05% sodium azide). Fc receptors were blocked with Human TruStain FcX (BioLegend, 422301) at 1:50 dilution for 5 minutes at 4°C. Then, cells were incubated for 30 minutes at 4°C in the dark with a staining solution containing CITE-seq antibodies (described below), BV421-labeled anti-CD3ε (clone SK7, BioLegend, 344833), and AF647-labeled CD19-tetramers (3 nM final concentration) for CAR T-cell phenotyping and detection. Subsequently, stained cells were conjugated with LIVE/DEAD Fixable Near-IR viability dye (Invitrogen, L34975) at 1:1000 dilution in PBS for 5 minutes at room temperature. Finally, cells were washed three times in cold cell media (RPMI, 10% FBS) before fluorescence-activated cell sorting (BD Biosciences, FACSAria Fusion). CAR^+^ sorting gates were drawn based on fluorescence of PBMCs from a similarly stained healthy donor.

Sorted endogenous T cells (CD3^+^CAR^-^) and CAR T cells (CD3^+^CAR^+^) were separately partitioned into droplets for single-cell RNA-seq/CITE-seq/TCR-seq via Chromium Next GEM Single-Cell 5’Kit v2 (10x Genomics, 1000263). RNA-seq libraries were prepared according to manufacturer protocols. CITE-seq libraries were prepared via the 5’ Feature Barcode Kit (10x Genomics, 1000256). TCR-seq libraries were prepared via the Chromium Single-Cell Human TCR Amplification Kit (10x Genomics, 1000252). All libraries (RNA-seq, CITE-seq, TCR-seq) were quantified via the Qubit dsDNA HS Assay Kit (Invitrogen, Q32851), quality-checked for fragment sizes via high-sensitivity D5000 screentapes (Agilent, 5067-5592), pooled, and sequenced (Illumina, Novaseq-6000 and Nextseq-550).

### CITE-seq antibody preparation

Thirty-one human “Cellular Indexing of Transcriptomes and Epitopes by Sequencing” (CITE-seq^62^) antibodies were obtained from BioLegend (TotalSeq-C reagents): anti-CD3ε (clone UCHT1, 300479), anti-CD5 (clone UCHT2, 300637), anti-TCRα/β (clone IP26, 306743), anti-TCRγ/δ (clone B1, 331231), anti-CD4 (clone SK3, 344651), anti-CD8α (clone SK1, 344753), anti-CD45RA (clone HI100, 304163), anti-CD45RO (clone UCHL1, 304259), anti-CCR7 (clone G043H7, 353251), anti-CD95 (clone DX2, 305651), anti-CD57 (clone QA17A04, 393321), anti-CD25 (clone BC96, 302649), anti-CD127 (clone A019D5, 351356), anti-CD103 (clone Ber-ACT8, 350233), anti-CXCR3 (clone G025H7, 353747), anti-CCR4 (clone L291H4, 359425), anti-CCR6 (clone G034E3, 353440), anti-PD-1 (clone EH12.2H7, 329963), anti-TIM-3 (clone F38-2E2, 345049), anti-LAG-3 (clone 11C3C65, 369335), anti-CD39 (clone A1, 328237), anti-TIGIT (clone A15153G, 372729), anti-CD27 (clone O323, 302853), anti-CD40L (clone 24-31, 310849), anti-GITR (clone 108-17, 371227), anti-OX40 (clone Ber-ACT35, 350035), anti-4-1BB (clone 4B4-1, 309839), anti-CD28 (clone CD28.2, 302963), mouse IgG1-κ isotype control (clone MOPC-21, 400187), mouse IgG2a-κ isotype control (clone MOPC-173, 400293), and mouse IgG2b-κ isotype control (clone MPC-11, 400381). Residual infusion product cells were stained with all 31 CITE-seq antibodies. Patient peripheral blood mononuclear cells were stained with all CITE-seq antibodies minus anti-CD3ε (30 in total) because T cells sorted from these samples were stained with BV421-labeled anti-CD3ε. To prepare the CITE-seq staining solution, the antibody pool was constructed and centrifuged at 14000 × *g* for 10 minutes at room temperature in FACS buffer (PBS, 2% BSA, 0.05% sodium azide) to remove aggregates. The antibody supernatant was extracted and diluted with fresh FACS buffer to appropriate staining concentrations.

### Analysis of CAR T-cell phenotype by flow cytometry

Cryopreserved biospecimens were thawed (RPMI, 10% FBS) and washed with cold FACS buffer (PBS, 2% BSA, 0.05% sodium azide). Fc receptors were blocked with Human TruStain FcX (BioLegend, 422301) at 1:50 dilution for 5 minutes at 4°C. Then, cells were incubated for 30 minutes at 4°C in the dark with a staining solution containing AF647-labeled CD19-tetramers (3 nM final concentration), BV421-labeled anti-CD3ε (clone SK7, BioLegend, 344833), AF488-labeled anti-CD8α (clone SK1, BioLegend, 344716), PE-labeled anti-CX3CR1 (clone 2A9-1, BioLegend, 341603), PE/Cy7-labeled anti-CD39 (clone A1, BioLegend, 328211), and BV605-labeled anti-CD57 (clone QA17A04, BioLegend, 393303). Subsequently, stained cells were conjugated with LIVE/DEAD Fixable Near-IR viability dye (Invitrogen, L34975) at 1:1000 dilution in PBS for 5 minutes at room temperature. Finally, cells were washed three times in cold FACS buffer (RPMI, 10% FBS) before flow cytometer analysis (BD Biosciences, FACSAria Fusion).

### Single-cell RNA-seq data processing

RNA-seq and paired CITE-seq reads were aligned to the GRCh38 reference genome, which was modified with the CAR transgene sequence used in axicabtagene ciloleucel, and quantified using the cellranger count (10x Genomics, version 7.0.0).^63^ Only filtered gene/feature-barcode matrices that contained barcodes with unique molecular identifier (UMI) counts that passed the quality control were used for downstream analyses.

### UMAP analysis and clustering on single-cell RNA-seq data

UMAP analysis and clustering were performed using the Seurat package (Version 4.3.0).^64^ Raw count matrices were first converted to Seurat objects before being further merged into one Seurat object, with protein expression added as the antibody-derived tag (ADT) assay. Cells with ˂300 genes detected or ˃10% mitochondrial RNA content were excluded from further analysis.

The raw count was log-normalized using the *NormalizeData* function with default options. The top 5,000 variable features were then identified using the *FindVariableFeatures* function with the default “vst” method. The data were centered and scaled using the *ScaleData* function, with additional regression against 1) the percent of mitochondrial RNA content and 2) difference in cell cycle S-phase score and G2M-phase score. Scaled data were then used as input for principal component analysis (PCA) based on variable genes using the *RunPCA* function. Data harmonization to remove patient-specific effects was performed on the principal components using the Harmony package (Version 1.0) through the *RunHarmony* function.^65^ Next, UMAP was constructed based on the first 30 harmony components. The same harmony components were used to construct the shared nearest neighbor (SNN) graph using the *FindNeighbors* function, which was then partitioned to identify clusters using the *FindClusters* function with default Louvain algorithm. These clusters were manually aggregated and classified as T-cell subsets based on known markers. CD8 and CD4 T-cell classifications were based on both gene and protein expression of CD4, CD8α, and CD8β.

For subset analysis, the corresponding subsets were extracted from the master Seurat object using the *subset* function. The above detailed preprocessing steps were repeated to generate the corresponding UMAP and subset annotations.

### Gene expression visualization

Gene expression was visualized in three ways: violin plots, heatmaps, and density plots. Violin plots were made based on normalized expression. Heatmaps were made based on average scaled expression, both using Seurat internal functions. Density plots were constructed using the Nebulosa package (version 1.6.0) to enhance visibility and mitigate sparsity of gene expression on UMAP. This is necessary for datasets with large number of cells (e.g. our CD8^+^ CAR-T dataset has ∼38k cells).

### Differential gene expression (DEG) analysis on single-cell RNA-seq data

DEG analyses were by default performed using the *FindMarkers* function in Seurat package, with default parameters and the appropriate ‘ident.1” and “ident.2” set as contrast. Unless otherwise stated, the results were filtered with p_val_adj < 0.05 and abs(avg_log2FC) > 0.25. Moreover, we also performed additional DEG analyses through a pseudo-bulk approach to better control for patient-specific effects, using the LibraDEG package (version 1.0.0)^66^ with default parameters (edgeR with LRT method). This was mainly performed when extracting timepoint-specific genes. Areas where this pseudo-bulk method is used are clearly stated in the figure legend.

### Timepoint-specific gene signature validation

To validate the timepoint-specific molecular signatures shared across patients and against external data, we adopted two independent and complementary approaches: 1) correlation of timepoint-specific gene expression, 2) label transfer from our data to external data.

For the correlation approach, we first conducted pseudo-bulk DEG analyses of T_exp_ vs T_per_ across all patients and kept all the statistically significant genes (padj <0.05) as timepoint-specific genes. Note that we did not filter by log fold-change to include even weak timepoint-specific signatures. Essentially, the goal here is to remove all non-timepoint-specific genes that may confound correlation analysis. The normalized gene expressions of these timepoint-specific genes were extracted from both our data and the external data. Following that, average expression was calculated for each sample (patient by timepoint). Correlations were then calculated pairwise for all samples.

For the label transfer approach, we adapted the Seurat standard data integration and label transfer workflow with default parameters. This is a complementary approach where, rather than at patient-level expression, individual cells in the external data and our data could be compared and similar timepoint-labels could be assigned.

### GSEA and pathway enrichment analysis

Gene Set Enrichment Analysis (GSEA) and pathway enrichment analysis were carried out using the clusterProfiler (version 4.4.4)^67^ package based on the msigdb database built in msigdbr package (version 7.5.1).^68^

### Exhaustion signatures from reference datasets

Besides standard msigdb exhaustion-related pathways, we obtained T-cell subset-specific signatures from Wherry et al. (see Supplementary Table 1 in their study).^53^ The reference study provides the most updated and comprehensive CD8^+^ (exhausted) T cell subsets, in the standard LCMV/Cl13 mouse models where T_ex_ was first defined. To refine the signature, we converted the mouse genes to their human orthologous genes and filtered the list to retain only the top 500 (by descending avg_log2_FC) statistically significant (p_val_adj < 0.05) genes for each subset.

### Regulon analysis

To construct regulons in a specific cell subset, we first extracted the raw count matrix of that subset. Then, we supplied it as input for the pySCENIC package (version 0.10.4) and ran through the workflow as detailed in its documentation and publication.^69,70^ More specifically, the base script was adapted from the “PBMC10K” example script on the pyScenic website.^71^ In addition, the arboreto component was used to speed up analysis. Each resultant regulon is a gene list with a central transcription factor and all its putative target genes determined through the SCENIC algorithm, and each set of regulons is sample/subset-specific and represents a putative regulatory unit in that specific sample/subset.

### Regulon or general gene set signature scoring

Individual cells were scored using the AUCell package (version 1.10.0)^69^ for a particular gene set from the msidb database or from a single-cell-derived regulon as follows. The normalized gene expression was first used as input into the *AUCell_buildRankings* function to score each cell for gene set enrichment and to build a ranking matrix. The signature score was then calculated as an AUC score using the *AUCell_calcAUC* function with all default parameters. In later revisions, Ucell package (version 2.0.1)^72^ was used in some re-analyses, as it is computationally more efficient and generates similar results as the AUCell package.

### Machine learning classification on regulon scores

To classify CD8^+^ CAR T cells at T_exp_ and T_per_, we first built a regulon set for each subset independently, assuming that regulatory networks for each timepoint may be different. Then, we merged all regulon sets into a master regulon set. We calculated the regulon scores using the master regulon set for all cells from the two subsets, resulting in a regulon score matrix. Each cell then served as an observation and each regulon as a feature for machine learning.

The machine learning classification models were built in the caret framework using the caret package (version 6.0-90).^73^ Briefly, we divided the input data into training (75%), validation (15%), and test sets (10%). We trained and optimized xgboost models^74^ through grid hyperparameter search and with a 5-fold cross-validation until reaching a best accuracy of >85% (refer to figures for specific accuracies).

We interrogated the final models for feature importance evaluation using the Shapley Additive Explanation (SHAP) method^75^ implemented in the SHAPforxgboost package (version 0.1.1).^76^ Average SHAPley values were used to rank the regulons. Top timepoint-determining regulons were visualized and further analyzed.

### Single-cell TCR-seq data processing

The TCR-seq reads from each sample were aligned to the 10x curated GRCh38 vdj reference genome and quantified using the cellranger vdj (10x Genomics, version 6.0.0). The results were then aggregated using the cellranger aggr function, with the source patient (donor) and timepoint (origin) of each sample supplied in the metadata to guide the clonotype assignment. The resultant clonotype and filtered contig annotation data were used for downstream analyses.

### Single-cell TCR-seq tracing and phenotype linking

Each unique clonotype is defined by the amino acid and nucleotide sequence of CDR3 regions for paired productive TCRα and TCRβ chains. TCR clonality overlap and tracing analyses were carried out using the immunarch package (version 0.9.0).^77^ Clones with the same TCR CDR3 sequences from the same patient at different timepoints were re-grouped and traced from infusion product to the post-infusion samples.

Phenotype-to-clonotype linking: To identify T_exp_ and T_per_ precursors, clonotypes among post-infusion T_exp_ and T_per_ cell populations were identified respectively and traced back to the infusion product cells. When linking the TCR clonotype to the corresponding cell phenotype, we keep only data with matching cell barcodes from both the TCR-seq and RNA-seq data.

Abundance tracing: Clonal abundance for each CAR T clone is calculated as the percentage of all CAR T cells occupied by a clone at a given timepoint. Each clone was assigned a phenotype at each timepoint based on the most frequently occurring phenotype. T_exp_ clones that were predominantly EM-exh had their abundances tracked at T_per1_/T_per2._ T_per1_/T_per2_ clones that were predominantly TE had their abundances tracked at T_exp_.

### CAR transgene mapping

The axicabtagene ciloleucel CAR design is documented.^78^ Its sequence was confirmed by Sanger sequencing of genomic DNA extracted from axicabtagene ciloleucel infusion products. The CAR sequence was then added to the GRCh38 FASTA and GTF files accordingly. A custom reference for cellranger was built from these annotation files using cellranger mkref (10x Genomics, version 7.0.0). The resultant custom reference was used for CAR transgene mapping through cellranger count.

### Data availability

All data are available from the authors upon reasonable request.

### Code availability

For reproducibility, code used for our analysis and figure-making is available at github (https://github.com/DanielGuoshuaiCao/TwoStageCART).

### Inclusion and Ethics Statement

Roles and responsibilities were agreed amongst collaborators in advance. Our findings are not expected to result in stigmatization, incrimination, or discrimination.

## ACKNOWLEDGEMENTS

We thank Phi Beta Psi, the Ullman Fund in Cancer Immunology, the Hoogland Lymphoma Research Pilot Projects, Chicago Immunoengineering Innovation Center, American Cancer Society Scholar Award (SG-22-136-01-1BCD) and NIH New Innovator award (1DP2AI144245) for financial support (to J.H.). Y.H. was supported by the University of Chicago MSTP Training Grant (T32GM007281). T.P. was supported by the University of Chicago CSTR training grant (T32HL007381). N.A. was supported by the University of Chicago MTCR training grant (T32CA009594). This project was supported by the National Center for Advancing Translational Sciences of the National Institutes of Health through Grant Number 5UL1TR002389-02 that funds the Institute for Translational Medicine. We thank the UChicago Human Immunologic Monitoring Facility for patient biospecimen cryopreservation, as well as the Pritzker School of Molecular Engineering Single-Cell Immunophenotyping Core and UChicago Genomics Facility for instrumentation to generate single-cell sequencing data. This work was completed in part with resources provided by the University of Chicago’s Research Computing Center. The authors thank Dr. Karen M. Watters for scientific editing of the manuscript.

## AUTHOR CONTRIBUTIONS

Y.H., G.C., J.P.K., and J.H. conceived and designed all experiments. J.H. supervised the project. T.A., P.A.R., and M.R.B. coordinated patient biospecimens for these studies. G.C. and T.P. performed all single-cell and machine-learning analyses. Y.H. organized execution of all experiments. Y.H. and E.T. sorted CAR T cells and generated single-cell data. G.C. and Y.H. jointly interpreted data. N.A. and Y.H. designed cartoons. E.T. performed flow cytometry validation. Y.H. and G.C. prepared the manuscript. J.H. and J.P.K. edited the manuscript. All authors reviewed the manuscript.

## COMPETING INTERESTS

P.A.R. reports Research Support/Funding: BMS, Kite Pharma, Inc./Gilead, MorphoSys, Calibr, Tessa Therapeutics, Fate Therapeutics, Xencor, and Novartis Pharmaceuticals Corporation. Speaker’s Bureau: Kite Pharma, Inc./Gilead; Consultancy on advisory boards: AbbVie, Novartis Pharmaceuticals Corporation, BMS, Janssen, BeiGene, Karyopharm Therapeutics Inc., Takeda Pharmaceutical Company, Kite Pharma, Inc./Gilead, Sana Biotechnology, Nektar Therapeutics, Nurix Therapeutics, Intellia Therapeutics, and Bayer. Honoraria: Novartis Pharmaceuticals Corporation. M.R.B. reports Membership on an Advisory Board or Consultancy for Kite/Gilead, Novartis, CRISPR Therapeutics, Autolus Therapeutics, BMS, Incyte, Sana Biotechnology, Iovance Biotherapeutics. He has served on a Speakers Bureau for BMS, Kite/Gilead, Agios, and Incyte. J.P.K. receives research support from Merck, Verastem, and iTeos; has served on a speaker’s bureau for Kite/Gilead; and has served on advisory boards for Verastem, Seattle Genetics, MorphoSys, and Karyopharm.

## Notes

### Summary of Updates

The statement regarding data accessibility was updated.

