## Supplementary Data for "Two-Stage CD8^+^ CAR T-Cell Differentiation in Patients with Large B-Cell Lymphoma"

**LIST OF SUPPLEMENTARY ITEMS**


- **Figure S1. Positron emission tomography/computed tomography (PET/CT) imaging for therapy response.**
- **Figure S2. CAR detection by CD19-tetramers.**
- **Figure S3. CAR T-cell population characteristics.**
- **Figure S4. CAR T-cell repertoire overlap and expansion dynamics analyses.**
- **Figure S5. Phenotypic heterogeneity of $CD8^+$ CAR T cells.**
- **Figure S6. Phenotypic heterogeneity of $CD4^+$ CAR T cells.**
- **Figure S7. Clonotypic heterogeneity of $CD8^+$ and $CD4^+$ CAR T-cell clusters.**
- **Figure S8. Clonotypic and phenotypic shifts among top 3000 clones.**
- **Figure S9. Timepoint-specific transcriptomic signatures.**
- **Figure S10. Characteristics of timepoint-determining regulatory networks.**
- **Figure S11. Expression of timepoint-determining regulatory networks.**
- **Figure S12. Phenotypic heterogeneity of infusion product $CD8^+$ CAR T cells.**
- **Figure S13. Characterization of exhausted-like EM $CD8^+$ CAR T cells.**

### SUPPLEMENTARY FIGURES AND LEGENDS

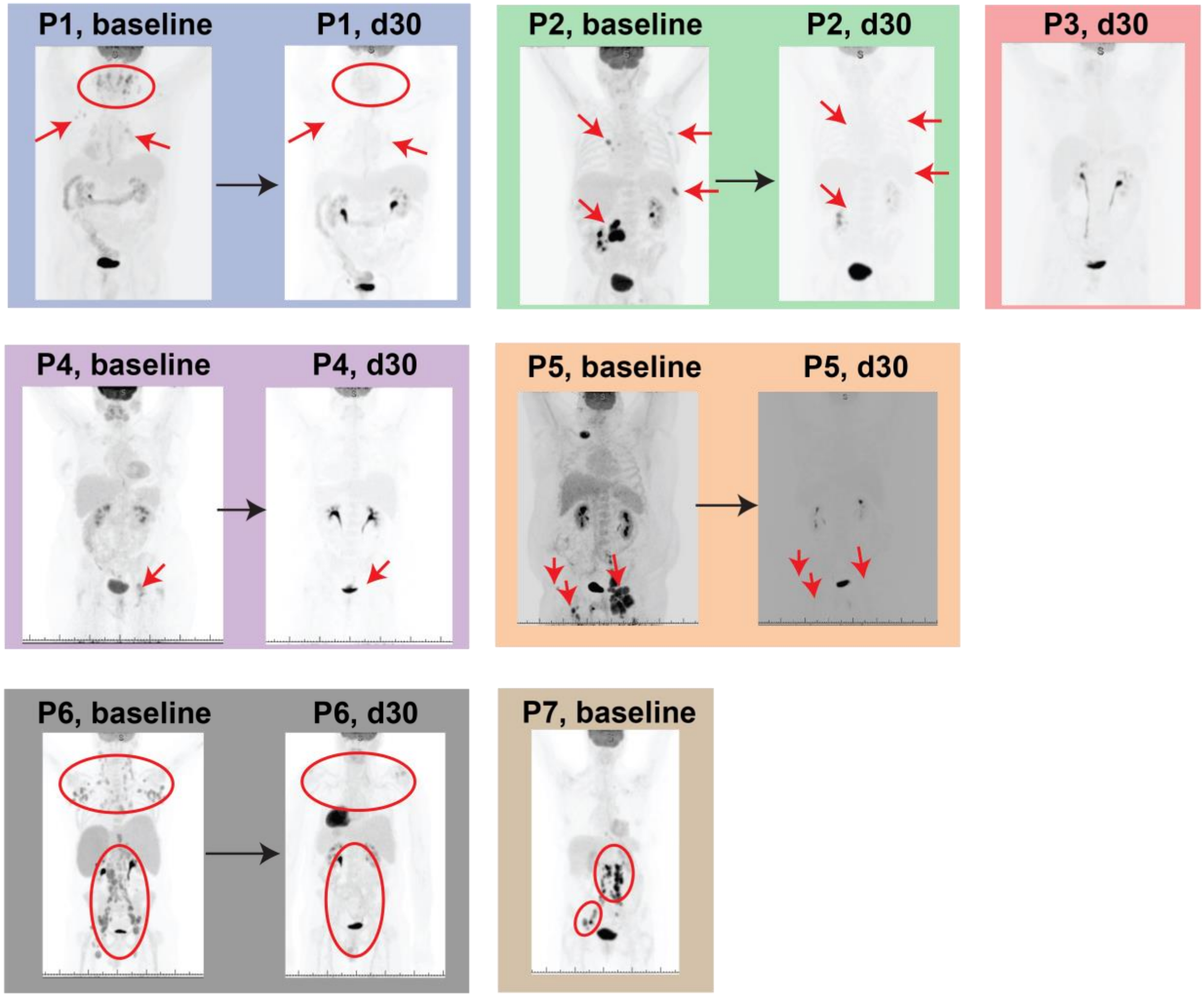


**Figure S1. Positron emission tomography/computed tomography (PET/CT) imaging for therapy response.** Baseline and 30-day response PET/CT scan assessment for patients (P1-7) with complete response. In all baseline images, major lymphoma locations are denoted by red arrows or circles. The same arrows and circles are displayed in the day 30 images, to indicate clearance of the tumor and complete response. Day 30 imaging for P7 was performed, but raw imaging data was not available.

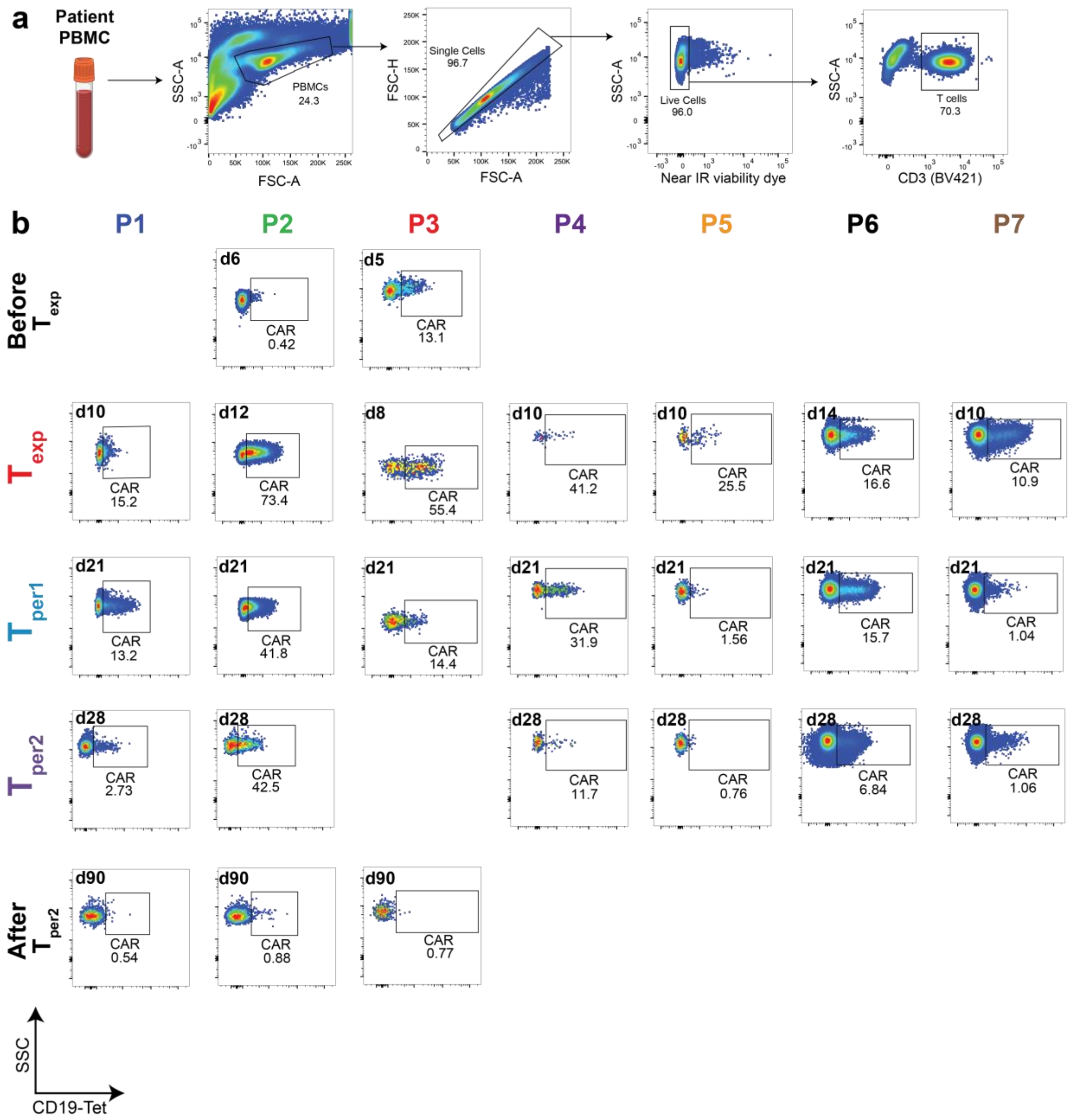


**Figure S2. CAR detection by CD19-tetramers.** Flow plots demonstrating use of Brilliant Violet 421-labeled anti-CD3ε and Alexa Fluor 647-labeled CD19-tetramers for CAR T-cell detection, quantification, and sorting. (**a**) Gating strategy for $CD3^+$ T cells from patient PBMCs. (**b**) Gating for CAR T cells. PBMCs from a healthy donor (biological control) were used to draw $CAR^+$ gates. PBMCs from each of the seven patients (labeled by columns) were analyzed at timepoints throughout the course of therapy (descending chronological order by rows).

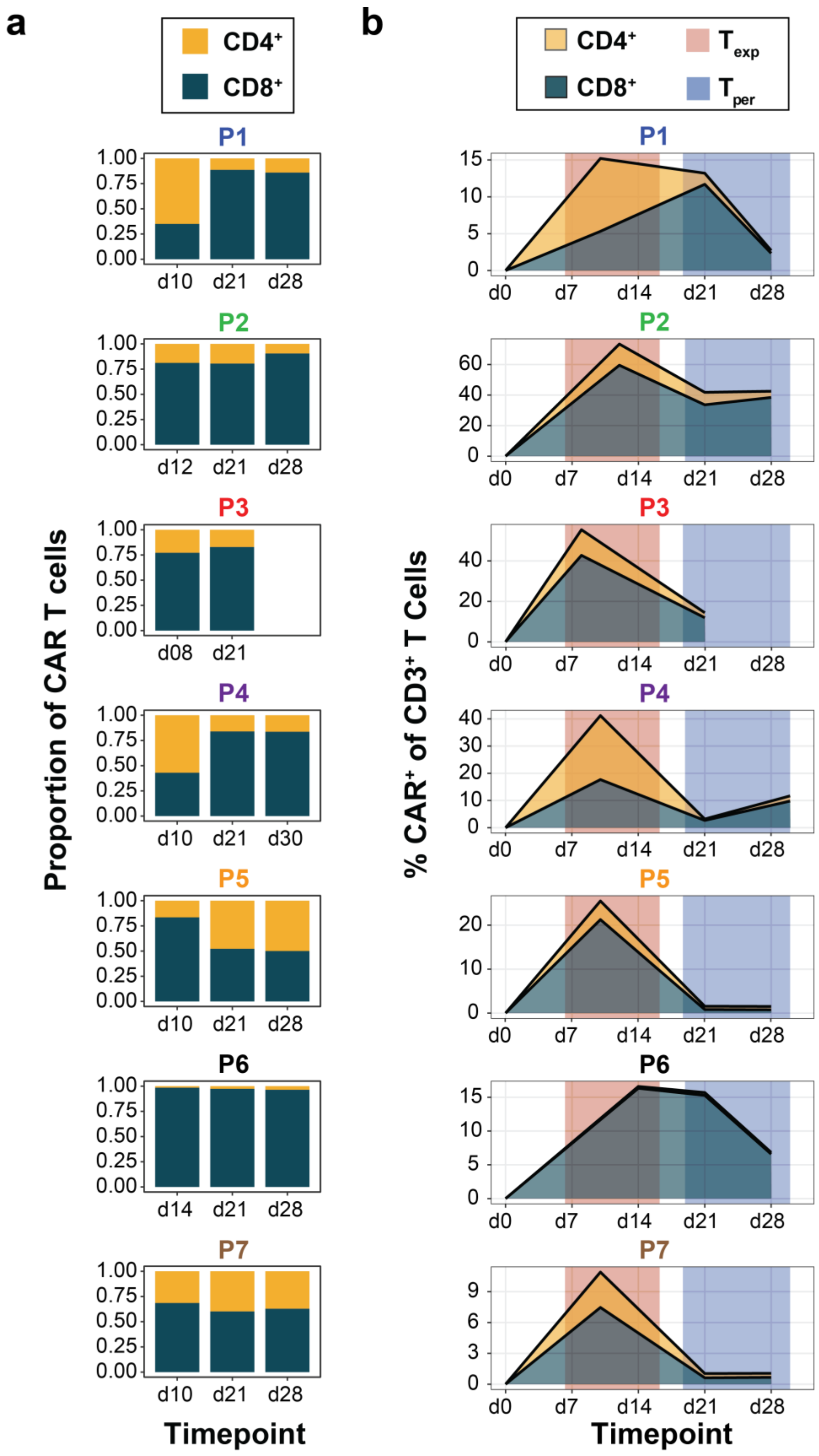

a
CD4+
CD8+
P1
P2
P3
P4
P5
P6
P7
Proportion of CAR T cells
Timepoint
b
CD4+
CD8+
Texp
Tper
% CAR+ of CD3+ T Cells
Timepoint

**Figure S3. CAR T-cell population characteristics.** (**a**) Stacked bar graphs indicating CD4 versus CD8 representation in CAR T cells for each patient at various timepoints. (**b**) Line plots describing the CAR T-cell population at various timepoints. Total CAR abundance (black line) is depicted for each patient. The area under this line is colored according to proportion of $CD4^+$ or $CD8^+$ CAR T cells. The $T_{exp}$ (peak expansion, day 8-14) and $T_{per}$ (post-peak persistence, days 21-28) timeframes are colored in the background accordingly.

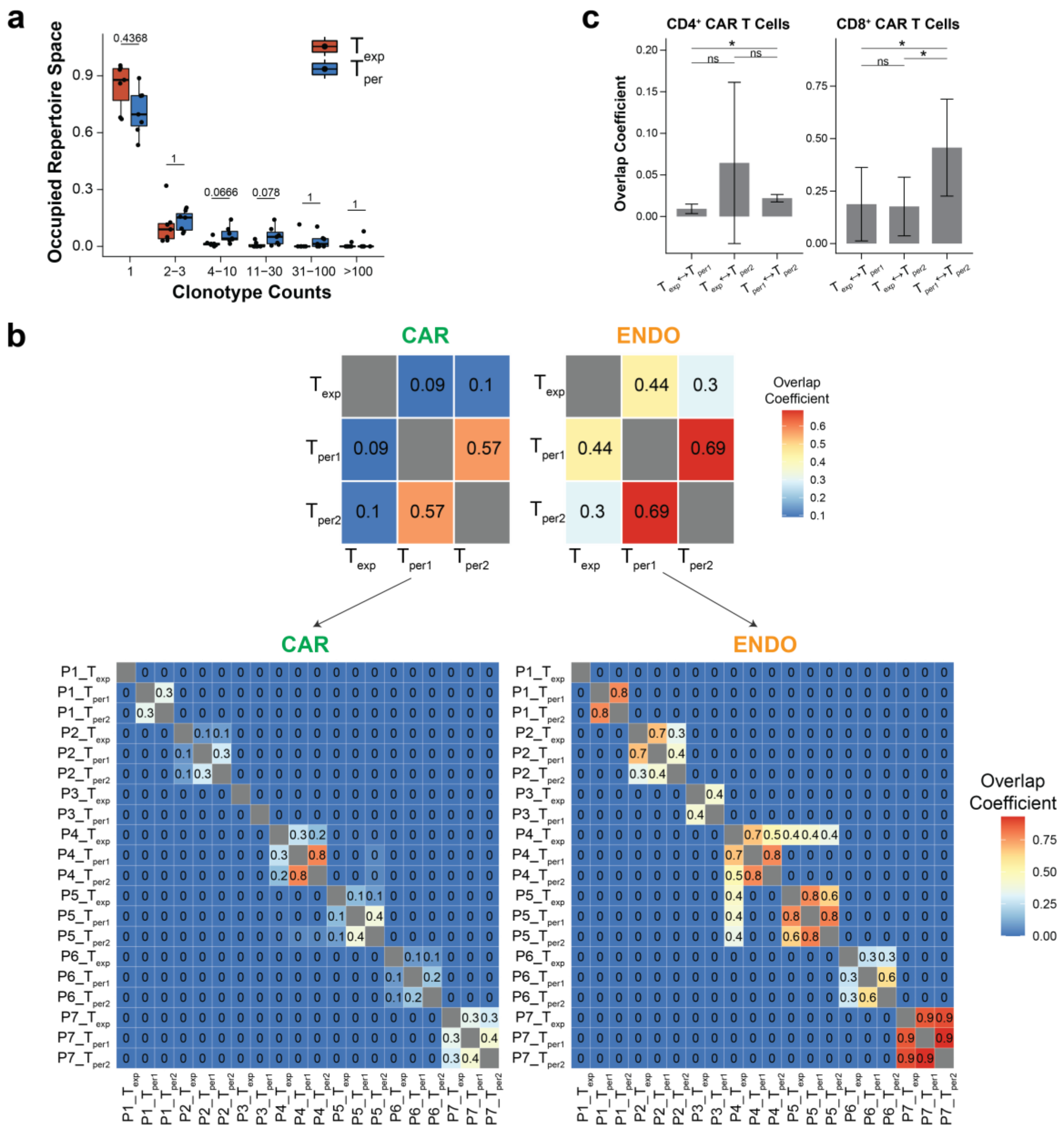


**Figure S4. CAR T-cell repertoire overlap and expansion dynamics analyses.** (**a**) Boxplot depicting the proportion of clonotypes of a given clone size. Proportions were compared between $T_{exp}$ (red) and $T_{per}$ (blue) by Wilcoxon Rank-Sum test, whereby *p*-values were labeled above each comparison. (**b**) Heatmap depicting overlap coefficients of TCR clonotypes from each timepoint (top) or sample (patient by timepoint, bottom) for sorted CAR ($CD3^{+}CAR^{+}$, left) and ENDO ($CD3^{+}CAR^{-}$, right) T cells. (**c**) Bar graphs depicting overlap coefficients comparing TCR clonotypes at $T_{exp}$, $T_{per1}$, and $T_{per2}$ for $CD4^{+}$ CAR T-cell (left) and $CD8^{+}$ CAR T-cell (right). Overlap coefficients were compared by *t*-test, whereby * indicates $p<0.05$ and ns indicates not significant.

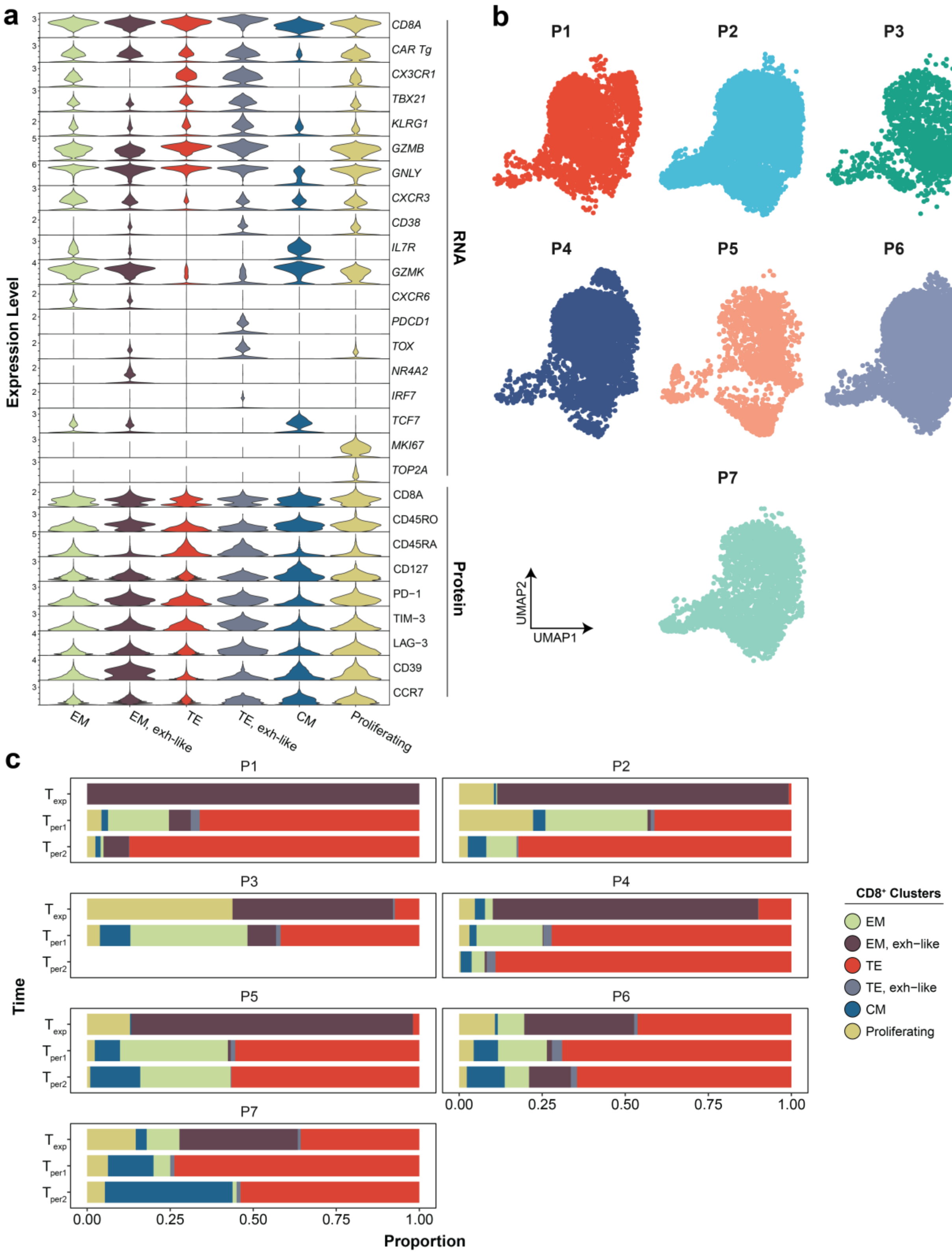

a
b
c
Expression Level
CD8A
CAR Tg
CX3CR1
TBX21
KLRG1
GZMB
GNLY
CXCR3
CD38
IL7R
GZMK
CXCR6
PDCD1
TOX
NR4A2
IRF7
TCF7
MKI67
TOP2A
CD8A
CD45RO
CD45RA
CD127
PD−1
TIM−3
LAG−3
CD39
CCR7
RNA
Protein
EM
EM, exh−like
TE
TE, exh−like
CM
Proliferating
P1
P2
P3
P4
P5
P6
P7
UMAP2
UMAP1
Time
$T_{exp}$
$T_{per1}$
$T_{per2}$
0.00
0.25
0.50
0.75
1.00
Proportion
CD8⁺ Clusters
EM
EM, exh−like
TE
TE, exh−like
CM
Proliferating

**Figure S5. Phenotypic heterogeneity of CD8$^+$ CAR T cells.** (**a**) Violin plots depicting normalized expression levels of key genes and proteins among the 6 CD8$^+$ CAR T-cell clusters. (**b**) Colored UMAPs depicting how CD8$^+$ CAR T cells from each patient are distributed on the overall harmonized UMAP. (**c**) Stacked bar graphs depicting proportions of CD8$^+$ CAR T cells represented in each cluster for each patient.

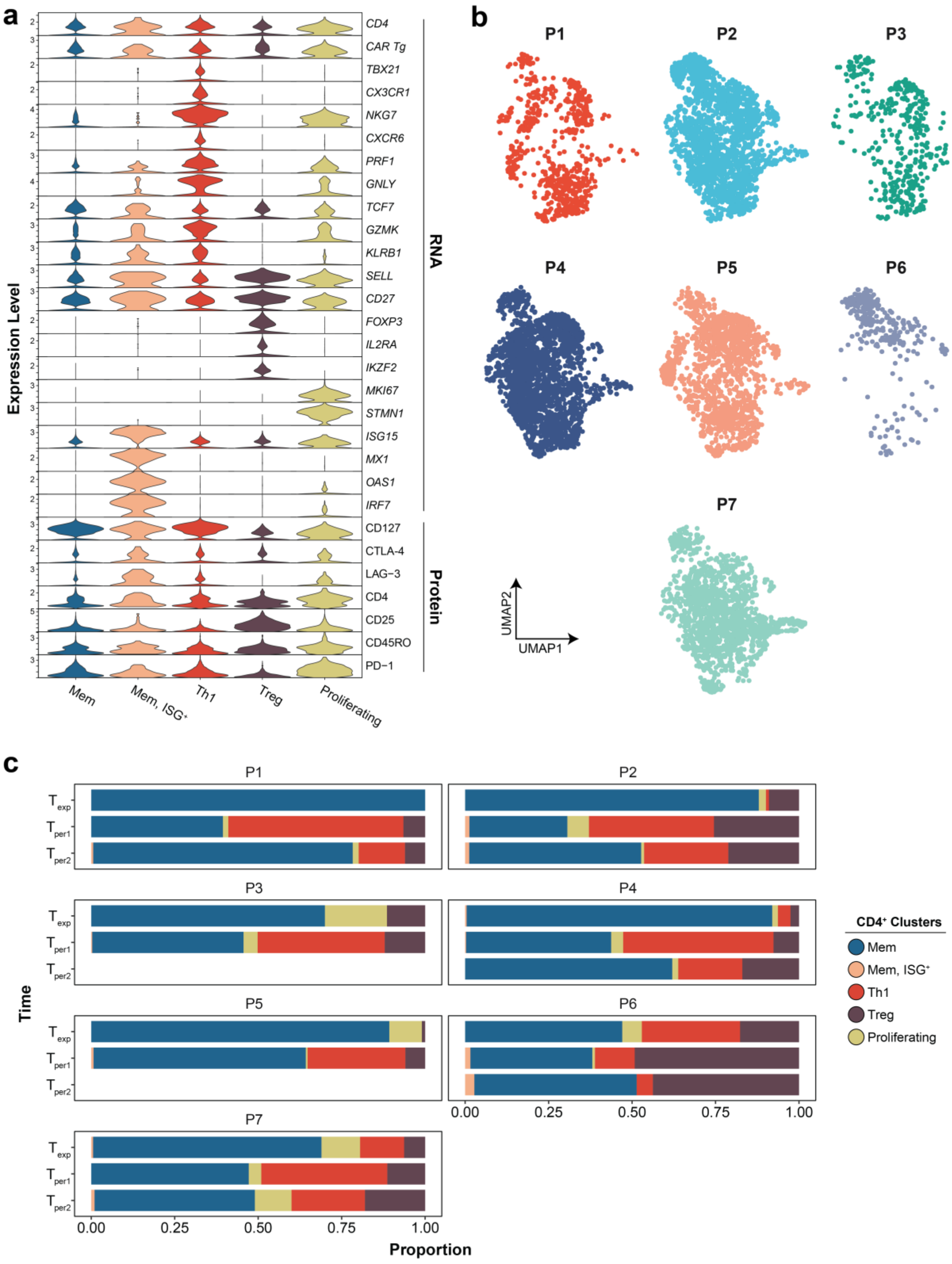
a
b
c
Expression Level
CD4
CAR Tg
TBX21
CX3CR1
NKG7
CXCR6
PRF1
GNLY
TCF7
GZMK
KLRB1
SELL
CD27
FOXP3
IL2RA
IKZF2
MKI67
STMN1
ISG15
MX1
OAS1
IRF7
CD127
CTLA-4
LAG-3
CD4
CD25
CD45RO
PD-1
RNA
Protein
Mem
Mem, ISG+
Th1
Treg
Proliferating
P1
P2
P3
P4
P5
P6
P7
UMAP2
UMAP1
Time
Texp
Tper1
Tper2
0.00
0.25
0.50
0.75
1.00
Proportion
CD4+ Clusters
Mem
Mem, ISG+
Th1
Treg
Proliferating

**Figure S6. Phenotypic heterogeneity of CD4$^+$ CAR T cells.** (**a**) Violin plots depicting normalized expression levels of key genes and proteins among the 5 CD4$^+$ CAR T-cell clusters. (**b**) Colored UMAPs depicting how CD4$^+$ CAR T cells from each patient are distributed on the overall harmonized UMAP. (**c**) Stacked bar graphs depicting proportions of CD4$^+$ CAR T cells represented in each cluster for each patient.

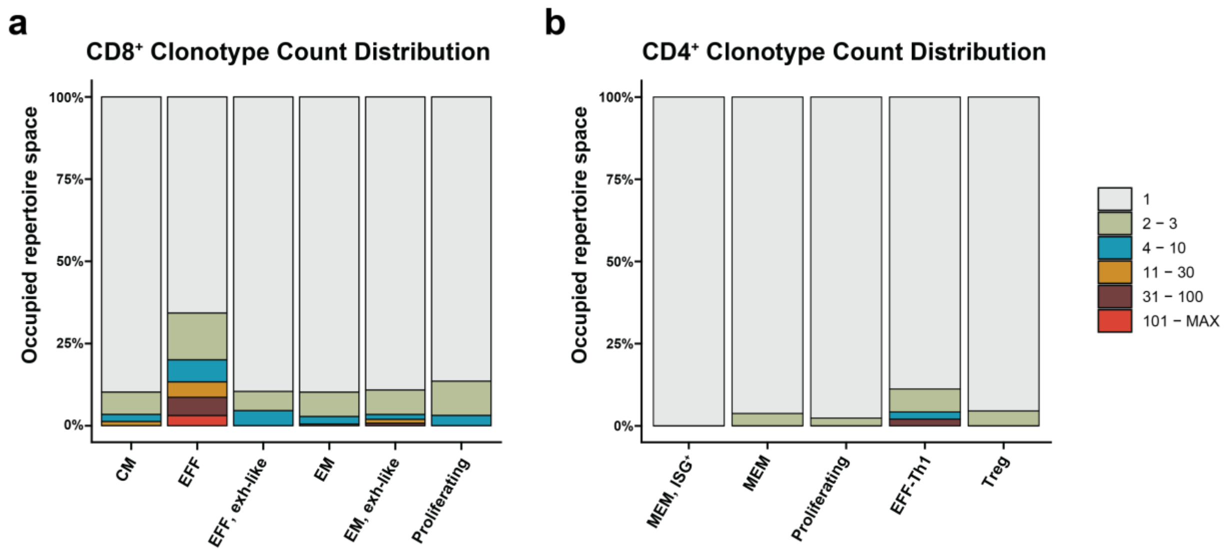


**Figure S7. Clonotypic heterogeneity of CD8⁺ and CD4⁺ CAR T-cell clusters.** Stacked bar graphs indicating the proportion of clonotypes of a given clone size in each CD8⁺ (**a**) and CD4⁺ (**b**) T-cell cluster. Most clonotypes had a clone size of 1.

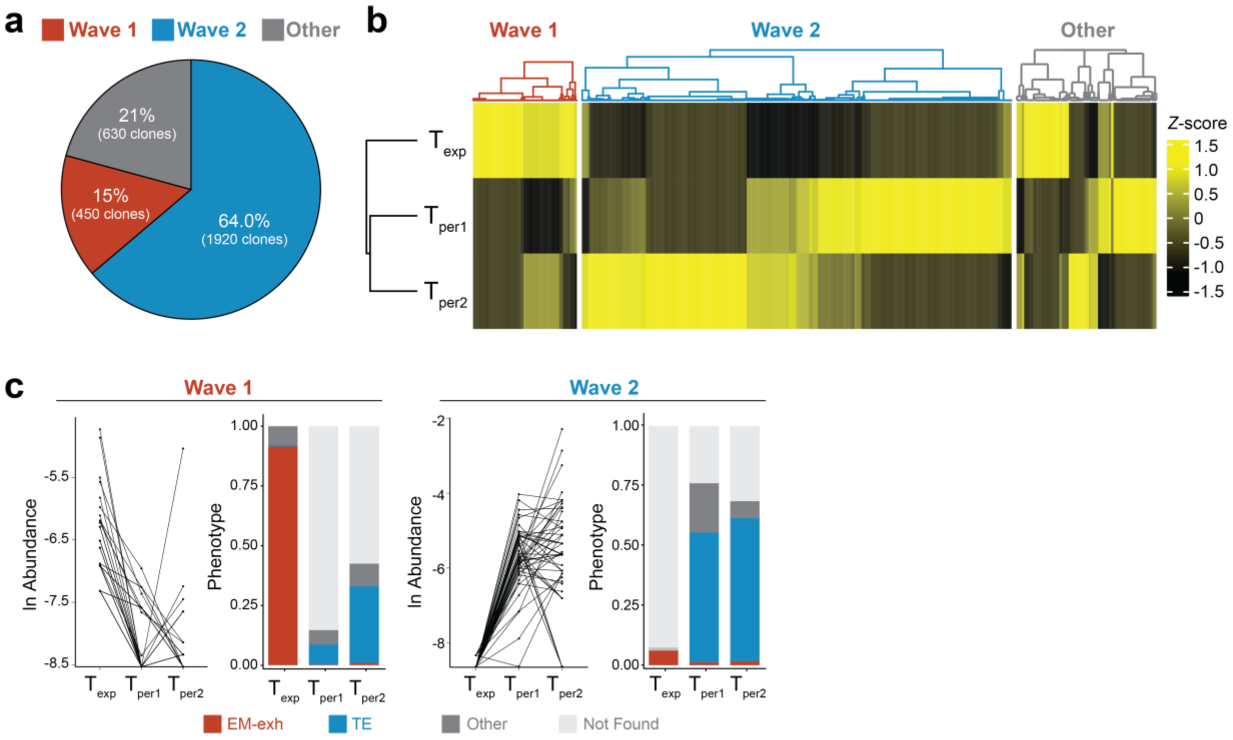


**Figure S8. Clonotypic and phenotypic shifts among top 3000 clones.** All figure panels are based on the clonotype-phenotype linked dataset (n=32432 cells). (**a**) Pie chart depicting proportion of the top 3000 clones classified as Wave 1, Wave 2, or Other. (**b**) Heatmap depicting normalized CAR abundance across timepoints for the top 3000 largest clones. (**c**) Clonal dynamics and phenotypic distribution for the top 3000 clones, grouped into Wave 1 and Wave 2. Left panels depict the change in clonal abundance for the top 30 (Wave 1) or top 50 (Wave 2) clones across timepoints, while right panels show the predominant phenotype distribution at each timepoint.

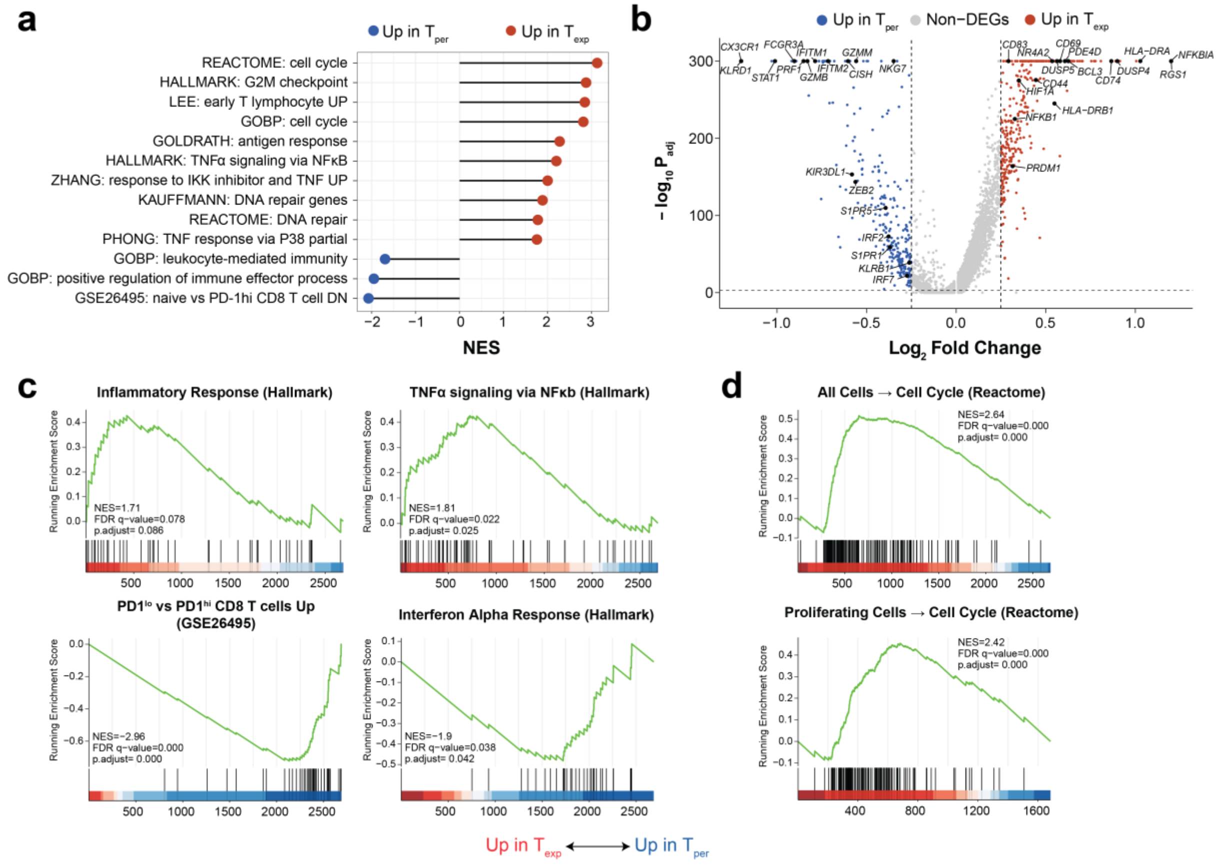


**Figure S9. Timepoint-specific transcriptomic signatures.** (**a**) Gene set enrichment analysis comparing $CD8^+$ CAR T cells at $T_{exp}$ and $T_{per}$ using the pseudobulk method. Gene sets were ordered by direction of upregulation and magnitude of enrichment. (**b**) Volcano plot depicting differentially expressed genes between $T_{exp}$ and $T_{per}$. Genes were colored according to direction of upregulation. (**c**, **d**) Enrichment plots for select gene sets related to T-cell and cytokine response (**c**) or proliferation (**d**). The proliferation gene set (**d**) was plotted for all $CD8^+$ CAR T cells (top) or the proliferating cell cluster only (bottom). Chosen gene sets are differentially expressed between $CD8^+$ CAR T cells at $T_{exp}$ and $T_{per}$.

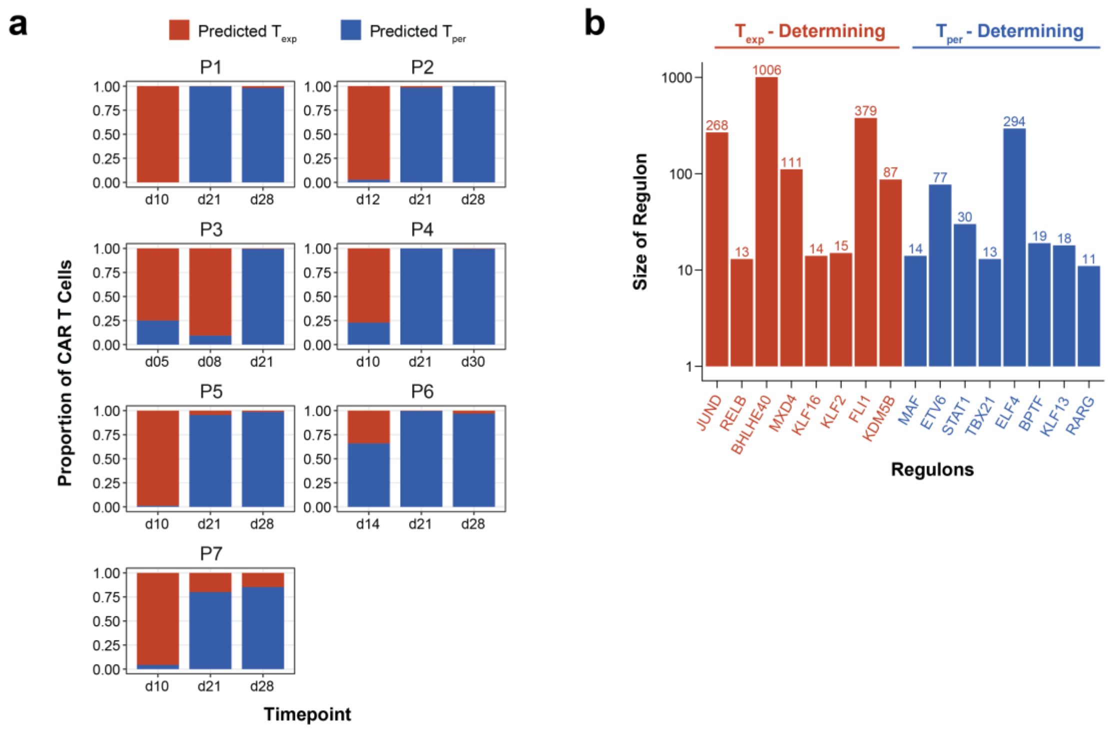


**Figure S10. Characteristics of timepoint-determining regulatory networks.** (**a**) Stacked bar graphs indicating the proportions of $CD8^+$ CAR T cells predicted to derive from the $T_{exp}$ or $T_{per}$ clonal expansion wave. Actual timepoints are labeled on the bottom of each graph. Predictions from the model were based on regulon expression transformed from single-cell transcriptomes. (**b**) Bar graphs depicting the number of genes in the top eight $T_{exp}$- or $T_{per}$-determining regulon.

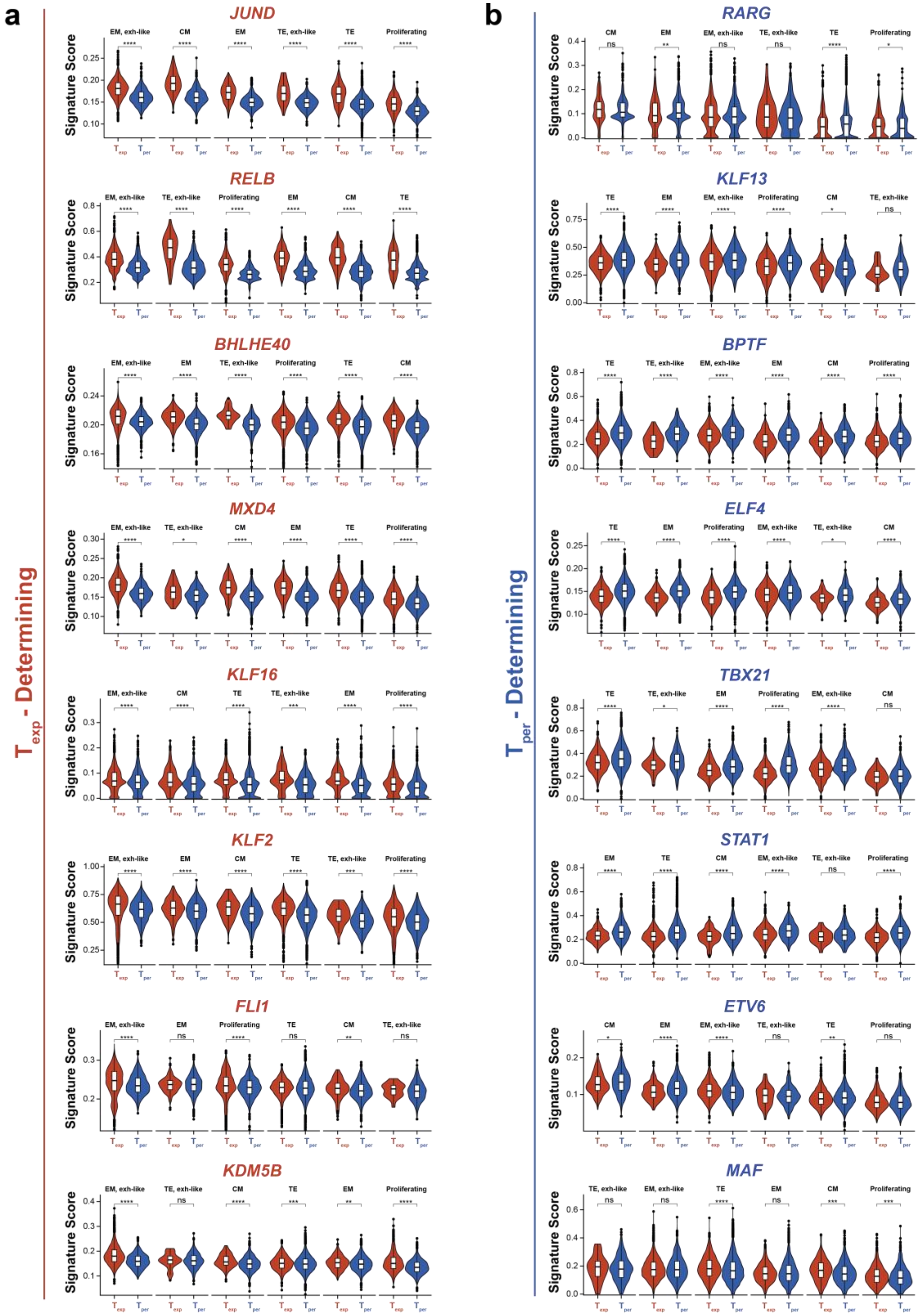

a
b
Texp - Determining
Tper - Determining
JUND
RELB
BHLHE40
MXD4
KLF16
KLF2
FLI1
KDM5B
RARG
KLF13
BPTF
ELF4
TBX21
STAT1
ETV6
MAF
Signature Score

**Figure S11. Expression of timepoint-determining regulatory networks.** Violin plots depicting expression of the top eight $T_{exp}$-determining (**a**) and $T_{per}$-determining (**b**) regulons among the six $CD8^{+}$ CAR T-cell clusters. Expression levels at $T_{exp}$ and $T_{per}$ were compared via Wilcoxon Rank-Sum test, whereby **** indicates $p<0.0001$, *** indicates $p<0.001$, ** indicates $p<0.01$, * indicates $p<0.05$, and ns indicates not significant.

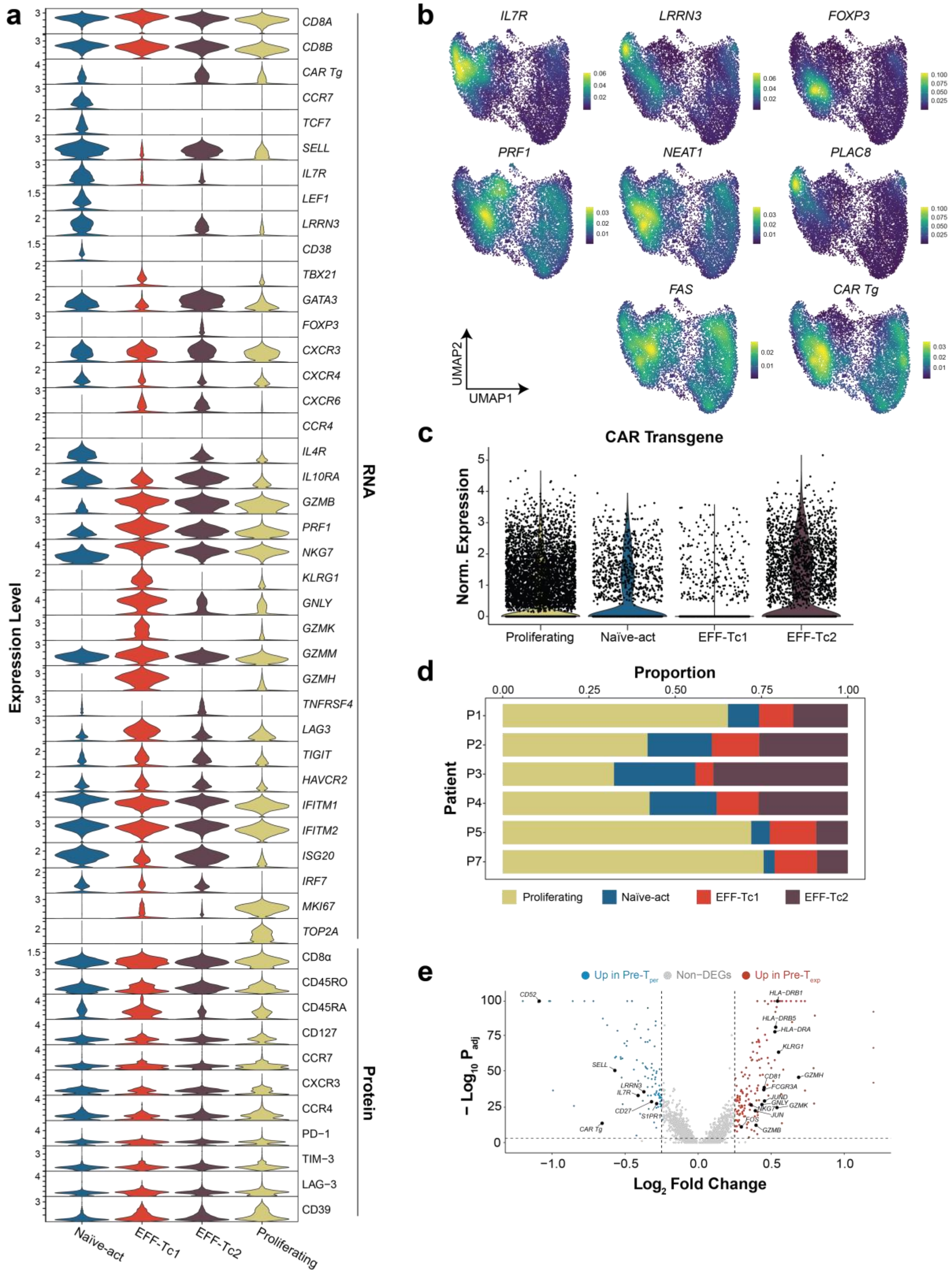
a
Expression Level
CD8A
CD8B
CAR Tg
CCR7
TCF7
SELL
IL7R
LEF1
LRRN3
CD38
TBX21
GATA3
FOXP3
CXCR3
CXCR4
CXCR6
CCR4
IL4R
IL10RA
GZMB
PRF1
NKG7
KLRG1
GNLY
GZMK
GZMM
GZMH
TNFRSF4
LAG3
TIGIT
HAVCR2
IFITM1
IFITM2
ISG20
IRF7
MKI67
TOP2A
RNA
CD8α
CD45RO
CD45RA
CD127
CCR7
CXCR3
CCR4
PD−1
TIM−3
LAG−3
CD39
Protein
Naïve-act
EFF-Tc1
EFF-Tc2
Proliferating
b
IL7R
LRRN3
FOXP3
PRF1
NEAT1
PLAC8
FAS
CAR Tg
UMAP1
UMAP2
c
CAR Transgene
Norm. Expression
Proliferating
Naïve-act
EFF-Tc1
EFF-Tc2
d
Proportion
0.00
0.25
0.50
0.75
1.00
Patient
P1
P2
P3
P4
P5
P7
Proliferating
Naïve-act
EFF-Tc1
EFF-Tc2
e
Up in Pre-T per
Non−DEGs
Up in Pre-T exp
− Log10 P adj
Log2 Fold Change
CD52
SELL
LRRN3
IL7R
CD27
S1PR1
CAR Tg
HLA−DRB1
HLA−DRB5
HLA−DRA
KLRG1
GZMH
CD81
FCGR3A
JUND
GNLY
GZMK
NKG7
JUN
FOS
GZMB

**Figure S12. Phenotypic heterogeneity of infusion product CD8$^+$ CAR T cells.** (**a**) Violin plots depicting normalized expression levels of key genes and proteins among the four CD8$^+$ infusion product clusters. (**b**) Density maps depicting expression density of key genes on the overall UMAP. (**c**) Violin plot depicting CAR transgene expression among the 4 clusters. (**d**) Stacked bar plots depicting distribution of each patient's infusion product CD8$^+$ T cells among the four clusters. (**e**) Volcano plot depicting differentially expressed genes between Pre-$T_{exp}$ and Pre-$T_{per}$. Genes were colored according to direction of upregulation.

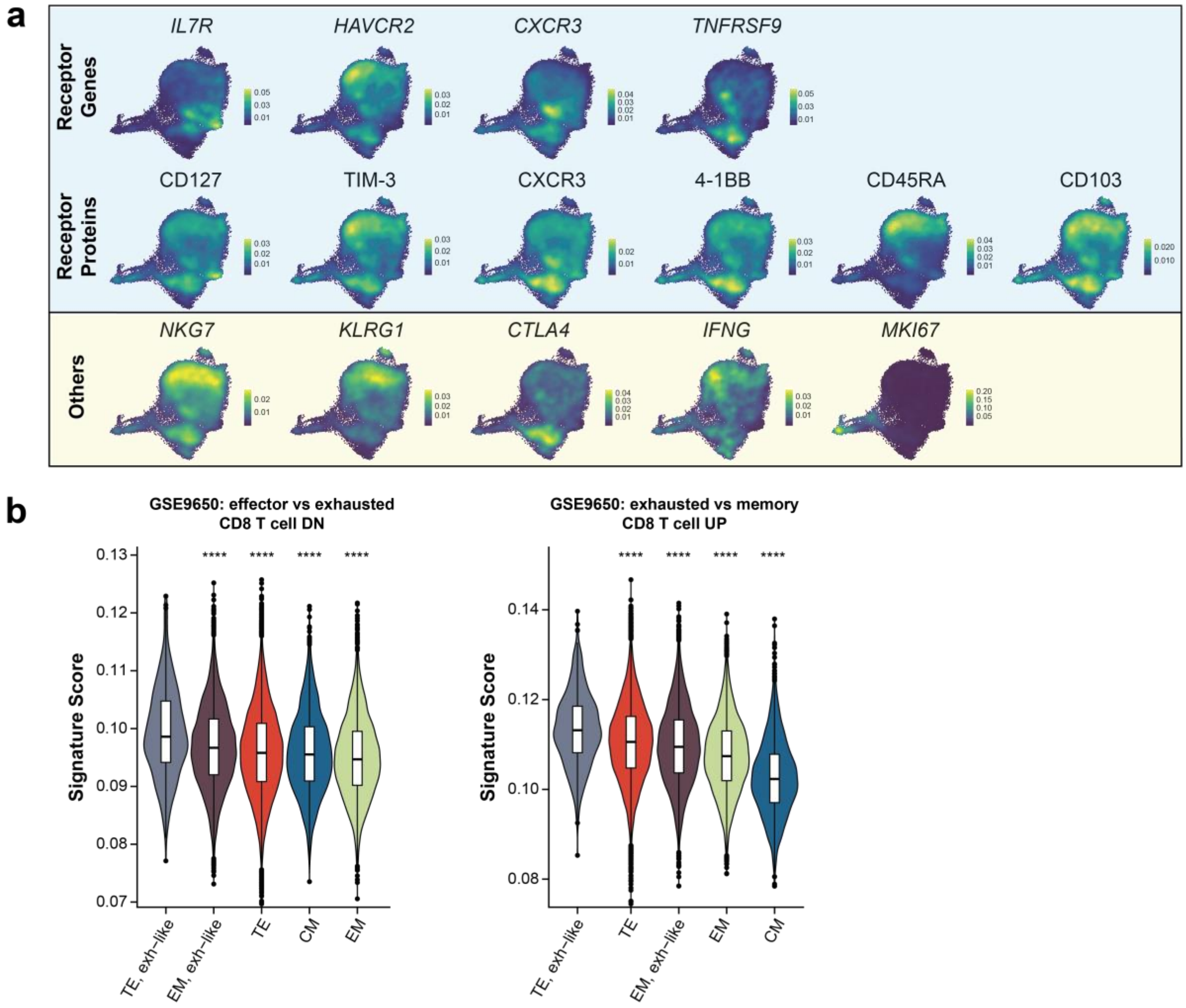


**Figure S13. Characterization of exhausted-like EM CD8+ CAR T cells.** (**a**) Density maps depicting expression levels of major T-cell genes and proteins, divided into categories. In the "receptor" category, proteins are placed directly beneath the corresponding gene. (**b**) Violin plots depicting expression of two exhaustion-associated gene sets, ordered by decreasing expression level per cluster. Expression levels were compared to that of the cluster with highest expression via Wilcoxon Rank-Sum test, with p-values adjusted for multiple hypotheses testing using the Benjamini-Hochberg method, whereby **** indicates p < 0.0001.
